# Rebalance the inhibitory system in the elderly brain: Influence of balance learning on GABAergic inhibition and functional connectivity

**DOI:** 10.1101/2024.09.16.613232

**Authors:** Xinyu Liu, Selin Scherrer, Sven Egger, Song-I Lim, Benedikt Lauber, Ileana Jelescu, Alessandra Griffa, Giulio Gambarota, Wolfgang Taube, Lijing Xin

**Affiliations:** Laboratory for functional and metabolic imaging (LIFMET), Ecole Polytechnique Fédérale de Lausanne, Lausanne, Switzerland; CIBM Center for Biomedical Imaging, Switzerland; Animal Imaging and Technology, Ecole Polytechnique Fédérale de Lausanne (EPFL), Lausanne, Switzerland; Institute of Physics, École Polytechnique Fédérale de Lausanne, Lausanne, Switzerland; Department of Neurosciences and Movement Science, University of Fribourg, Fribourg, Switzerland; Department of Radiology, Lausanne University Hospital, Lausanne, Switzerland; Medical Image Processing Laboratory, Neuro-X Institute, Ecole Polytechnique Fédérale de Lausanne (EPFL), Geneva, Switzerland; Leenaards Memory Center, Lausanne University Hospital and University of Lausanne, Lausanne, Switzerland; Faculty of Pharmacy, University of Rennes, Rennes, France

## Abstract

Aging involves complex processes that impact the structure, function, and metabolism of the human brain. Declines in both structural and functional integrity along with reduced local inhibitory tone in the motor areas, as indicated by reduced γ-Aminobutyric acid (GABA) levels, are often associated with compromised motor performance in elderly adults. Using multi-modal neuroimaging techniques including magnetic resonance spectroscopy (MRS), diffusion magnetic resonance imaging (MRI), functional MRI as well as transcranial magnetic stimulation to assess short interval intracortical inhibition (SICI), this study explores whether these age-related changes can be mitigated by motor learning. The investigation focused on the effects of long-term balance learning (3 months) on intracortical inhibition, metabolism, structural and functional connectivity in the cortical sensorimotor network among an elderly cohort. We found that after three months of balance learning, subjects significantly improved balance performance, upregulated sensorimotor cortical GABA levels and ventral sensorimotor network functional connectivity (VSN-FC) compared to a passive control group. Furthermore, correlation analysis suggested a positive association between baseline VSN-FC and balance performance, between baseline VSN-FC and SICI, and between improvements in balance performance and upregulation in SICI in the training group, though these correlations did not survive the false discovery rate correction. These findings demonstrate that balance learning has the potential to counteract aging-related decline in functional connectivity and cortical inhibition on the ‘tonic’ (MRS) and ‘functional’ (SICI) level and shed new light on the close interplay between the GABAergic system, functional connectivity, and behavior.

## Introduction

Aging is accompanied by progressively declined motor functions which can be linked to a broad range of neurochemical, structural and functional changes in the human brain [1]. Of particular relevance are aging-related deficits in cortical inhibitory functions, which in turn have been shown to be associated with various aspects of declined motor functions [2]. At the cortical level, motor inhibition is largely mediated by γ-Aminobutyric acid (GABA) and its associated GABAergic processes. As the major inhibitory neurotransmitter in the brain, GABA is thought to be crucial in maintaining a balance between excitation and inhibition within the central nervous system to facilitate efficient motor control and function. For example, it has been shown that a decline of sensorimotor cortical GABA level in elderly is linked to motor function deficits [3], [4]. Deficiencies or abnormalities in GABAergic activity have also been documented in cognitive and/or movement disorders among elderly population, such as Alzheimer’s disease [5] and Parkinson’s disease [6].

In addition to the neurochemical changes, aging was also shown to be paralleled by widespread functional and structural brain changes. Studies have consistently shown that aging is accompanied by declined resting-state functional connectivity within the sensorimotor network, which is associated with motor performance deficits [4], [7]. In addition, several studies indicated close links between the cortical GABAergic inhibitory system and functional connectivity [8], [9], [10], [11]. Structural MRI studies have found that motor performance declines in the elderly can be related to grey matter atrophy in motor cortical regions [12]. Taken together, these findings suggest that aging-related brain alterations, particularly in the cortical sensorimotor system, are crucial in motor performance decline among elderly adults.

An important question regarding the described aging-related brain changes is whether they can be counteracted by motor learning interventions. In young subjects, neuroimaging techniques have been used to investigate the neuroplastic mechanisms underlying the acquisition and adaptation of motor skills such as juggling [13], cycling [14] and golfing [15]. These studies have unveiled extensive structural and functional adaptations across the brain in response to motor learning. One particularly promising motor learning paradigm is balance learning, where participants practice balance tasks to enhance their ability to maintain postural stability. Previous studies have reported that balance learning resulted in grey matter changes in cerebellum, hippocampus and frontal regions as well as cortical motor areas [16], [17]. Furthermore, changes in functional connectivity in fronto-parietal networks, sensorimotor networks and the striatum were also reported in response to balance learning [16], [18], [19]. With respect to the aim of the present study to better understand the role and adaptation of inhibitory mechanisms in the cortex, learning of challenging balance tasks over several weeks has been demonstrated to increase short-interval intracortical inhibition (SICI), indicating upregulation of GABAergic processes [20], [21]. However, it is important to highlight that *first*: the majority of these studies focused on a younger population, leaving it unclear whether these interventions could elicit similar plasticity effects in elderly adults; and *second*: the question of how structural, functional and metabolic changes induced by balance learning interact with each other has not yet been systematically addressed. In one study including only young adults [22], it was reported that 6 weeks of juggling practice resulted in increased sensorimotor network functional connectivity (SNFC). Moreover, these changes were associated with modulation in the local GABA concentration levels in the primary motor cortex (M1) and with changes in motor performance. This suggests that alterations in resting-state network strength following motor learning are, at least in part, related to GABAergic processes.

In this study, we aimed to examine brain plasticity in response to long-term balance learning among an elderly cohort in comparison to a passive control group. Using a multi-modal approach, we seek to further disentangle associations between plastic changes that are usually assessed in separate studies due to the complexity of combining state-of-the-art imaging (MRS, MRI, fMRI) with neurophysiological methods. The primary outcome of the study is the training effects on cortical GABA level, SICI level and balance performance. Based on previous reports showing lower sensorimotor cortical GABA and functional connectivity levels in elderly adults [4], [7], [23] but upregulation of SICI after balance learning [20], we hypothesized that long-term balance learning would lead to enhanced sensorimotor cortical inhibition and SNFC. We further hypothesized that the modulation in cortical inhibition would be related to changes in behavior (i.e., balance performance). In addition, potential associations between the level of intracortical inhibition/GABA on one hand and SNFC and balance performance on the other hand were investigated both during the pre-measurement and for delta values (post minus pre). This was considered to be important in order to establish the influence of GABAergic processes on functional coupling of brain regions (i.e., functional connectivity) and motor performance (i.e., balancing). We also investigated whether there are concomitant alterations in the anatomical projections (white matter pathways) in the sensorimotor network.

Our study employed a multi-modal approach combining 7 T diffusion MRI (dMRI), functional MRI (fMRI), ^1^H-MRS and a paired-pulse paradigm with transcranial magnetic stimulation (TMS). Specifically, we sought to investigate the effects of three months of balance learning on both structural and functional connectivity within the cortical sensorimotor network, as well as the GABAergic processes within the sensorimotor cortex, a key node within the network. Due to its established importance in motor inhibition and functioning [24], GABA was the main metabolite of interest which can be reliably quantified using editing-based ^1^H-MRS at 7 T [25]. Furthermore, we also conducted an explorative analysis on other metabolites using short echo-time MRS. Complementary to MRS which is capable to detect the local total concentration of GABA, TMS was used to specifically detect the activation of GABA_A_ receptors during balancing in the primary motor cortex. Thus, TMS can more accurately quantify the degree of cortical inhibition, which involves GABA acting on synaptic GABA_A_ receptors [26], while participants are actually executing the trained motor task. The use of these two complementary methods in an interventional study is important to better understand the relationship between TMS measures of cortical inhibition reflecting synaptic GABAergic phasic inhibitory activity and MRS measurements reflecting total concentration of GABA [27].

## Methods

### Study design and participants

Forty participants aged 64 – 82 years old (18 males, 22 females) were recruited in this study. The participants were randomly assigned to a balance learning (20 participants) or a control group (20 participants). The sample size was chosen based on a power analysis conducted prior to the study which can be found in the **Supplementary Materials**, “sample size calculation” section. All participants were cognitive unimpaired and free of any known neurological and orthopedic diseases or injuries as well as contradictions to MRI and TMS measurements. Two participants in the learning group and two participants in the control group did not complete the post-measurement, thus only their data from the pre-measurement were used for analysis. Two participants in the learning group and one in control group did not participate in the MR examinations thus only data for balance learning and SICI were used for analysis; two participants in the learning group and two in the control group did not have SICI and balance performance data acquired successfully. Detailed demographics of participants can be found in **Table 1**. All participants provided written informed consent prior to the experiments and the experiments were performed with the approval of the local ethics committee (Commission cantonale d’éthique de la recherche sur l’être humain (CER-VD); ID: ID 2021-00378). The study consisted of two (PRE and POST) identical MR, TMS and balance measurements three months apart. Between PRE and POST measurements, participants in the training group engaged in three months of progressive, multifaceted balance learning. TMS and balance performance measurements were performed on the same day within a week after the end of training, MR measurements were performed within two weeks after the end of training.

**Table 1.** Demographics and MRS data quality parameters of participants. SNR for MEGA-sSPECIAL and sSPECIAL is calculated using height of GABA or NAA peak divided by noise standard deviation between 9.5 to 10 ppm, respectively. MRS quality parameters are calculated only for participants who participated in MR measurements. No significant difference was found between pre and post measurements within group or between two groups (p > 0.05).

|  |  | Number of<br>participants | Age | Sex | MEGA-sSPECIAL<br>SNR | sSPECIAL<br>linewidth | sSPECIAL<br>SNR |
| --- | --- | --- | --- | --- | --- | --- | --- |
| Balance<br>learning<br>group | Pre | 20 | 70.9 ±<br>4.5 | 9M/11F | 9.9 ± 3.0 | 12.8 ± 2.4 | 434.7 ±<br>103.6 |
|  | Post | 18 | 71.2 ±<br>4.0 | 8M/10F | 10.8 ± 3.1 | 11.6 ± 2.3 | 433.2 ±<br>105.8 |
| Control<br>group | Pre | 20 | 70.2 ±<br>4.7 | 9M/11F | 7.4 ± 2.27 | 11.6 ± 2.0 | 393.9 ± 91.5 |
|  | Post | 18 | 69.6 ±<br>4.6 | 8M/10F | 8.8 ± 2.6 | 11.3 ± 3.3 | 397.2 ± 76.1 |

### Balance learning

Participants in the training group were trained 3 times per week (a total of at least 30 training sessions) for 45 minutes in supervised group sessions. Each session consisted of different balancing exercises for the lower extremities, such as standing with one leg on mobile, uneven, or thin surfaces. These exercises could be adapted, making them easier with some support at the beginning or making them more difficult by reducing the base of support or adding a secondary task for the upper extremities, for example. The basic goal for the participants was to stabilize their posture under increasingly challenging conditions. The sessions started with a warm-up and finished with a cool down. A detailed balance learning plan is provided in the supplementary material. Participants in the control group were instructed to maintain their regular lifestyle without engaging in any new training modality.

### Balance performance measurement

Balance performance was evaluated using wobble boards, on which participants stood with both feet and hands held akimbo. The devices have a rounded bottom, creating an unstable surface. There were four wobble boards with different levels of difficulty by decreasing the base of support. The smaller the support surface the more difficult was the balance task. Participants were instructed to remain as stable as possible for twenty seconds while visually fixing a cross taped to the wall. Trials counted as successful if the participants did not use any external assistance and maintained balance on the board. Participants began with two trials on the easiest device. If they succeeded in at least one out of the two trials, they moved on to the next level of difficulty (i.e., wobble board with smaller base of support) again having two trials. Before a familiarization trial, participants were given time to locate their preferred position at the start of each new level. The pause between trials was approximately 1 minute. The wobble boards were placed on a force plate (508 × 464 mm; OR 6 – 7 force platform; Advanced Mechanical Technology Inc., Watertown, MA, USA) to record the center of pressure movements. The force plate signals were sampled at a frequency of 4 kHz and amplified 4000 times (GEN 5, Advanced Mechanical Technology Inc., Watertown, MA, USA). Recordings were analyzed offline with Matlab (R2021b, The MathWorks, Inc., Natick, MA, USA). The mean sway areas in cm^2^ (area of smallest ellipse that covers 95% of center of pressure data points) of the two attempts on the highest wobble board level that was achieved at pre-and post-measurement, ensuring performance comparisons between the same wobble board level within the same participant, were taken as the balance performance parameter. Thus, smaller sway area indicates better balance performance. There was at least a break of 2 days, at most one week between the last balance training session and the post-test.

### MR Data acquisition

All MR measurements were performed on a 7-T/68-cm MR scanner (Magnetom, Siemens Medical Solutions, Erlangen, Germany) equipped with 80 mT/m gradients and with a single-channel quadrature transmitter and a 32-channel receiver coil (Nova Medical Inc., MA, USA).

A high-resolution MP2RAGE [28] sequence was used to acquire structural brain image and for MRS voxel placement (voxel size = 0.6 × 0.6 × 0.6 mm^3^, TR/TE = 6000/4.94 ms, TI1/TI2 = 800/2700 ms, slice thickness = 0.6 mm, field of view = 192 × 192 × 154 mm^3^, acquisition time = 10 min 8 s).

GABA concentrations in the left sensorimotor cortex were assessed by edited single-voxel spectroscopy acquired by the MEGA-sSPECIAL (TR/TE = 4000/80 ms) sequence [25] with the following parameters: average number = 128, number of datapoints = 2048, spectral width = 4000 Hz, volume of interest = 30 × 20 × 20 mm^3^, acquisition time = 8 min 32 s. The sequence used a broad band asymmetric adiabatic inversion pulse with 500 Hz of inversion bandwidth applied at 1.9 ppm (edit-on) and highly selective Gaussian pulse with 98 Hz of inversion bandwidth applied at 1.7 ppm (edit-off) to suppress co-edited macromolecule signals. A short echo-time sSPECIAL sequence (TR/TE = 4000/16 ms) was used for metabolite profiling for a number of other neuro-metabolites (average number = 100, number of datapoints = 2048, spectral width = 4000 Hz, volume of interest = 30 × 20 × 20 mm^3^). A dielectric pad was positioned near the sensorimotor cortex to improve the transmit field efficiency [29]. The MRS voxel was placed in the left precentral gyrus according to our previous fMRI study which assessed brain activation during observation and motor imagery of different balance tasks [30].

Resting-state functional MRI images were acquired using a 2D multi-slice gradient-echo echoplanar imaging (EPI) sequence (TR = 1550 ms, TE = 26 ms, field of view = 198 × 198 mm^2^, in-plane resolution = 1.3 × 1.3 mm^2^, slice thickness = 1.3 mm, 100 slices, GRAPPA factor = 3, acquisition time = 6 min 52 s) [31].

Diffusion-weighted MRI images were acquired using a 2D multi-slice pulsed-gradient spin-echo (PGSE) EPI sequence (TR = 5000 ms, TE = 59.8 ms, b-values = 0 (4 repetitions), 200 (6 directions), 1000 (30 directions), 2000 (60 directions) s/mm^2^, voxel size = 2.0 × 2.0 × 2.0 mm^3^, field of view = 230 × 230 × 120 mm^3^, multi-band acceleration factor = 2, GRAPPA factor = 3, partial Fourier = 6/8, acquisition time = 8 min 45 s).

### MRS data processing and analysis

MR spectra were averaged after frequency drift and phase correction using FID-A [32] and analyzed by LCModel [33] for quantification. The MEGA difference spectra were fitted with a simulated basis set of five metabolites, including GABA, Glutamate (Glu), Glutamine (Gln), N-Acetylaspartic acid (NAA), and N-acetylaspartylglutamate (NAAG). The spectral range for LCModel analysis was set from 1.8 to 4.2 ppm except the nuisance signal in the range of 2.2 to 2.8 ppm. For the sSPECIAL spectra, an experimentally measured macromolecule spectrum and simulated spectra of the following 21 metabolites were included in the basis set for LCModel analysis: alanine (Ala), ascorbate (Asc), aspartate (Asp), Creatine (Cr), GABA, glucose (Glc), Gln, Glu, glycerophosphorylcholine (GPC), glycine (Gly), glutathione (GSH), lactate (Lac), myo-inositol (mIns), NAA, NAAG, phosphocreatine (PCr), phosphorylcholine (PCho), phosphorylethanolamine (PE), scyllo-inositol (sIns), serine (Ser), and taurine (Tau). All metabolites with Cramér–Rao lower bound (CRLB) larger than 30% were considered as non-detected. Only metabolites with mean CRLB smaller than 30% were included for further analysis.

Unsuppressed water signal was used as an internal reference for absolute quantification. To account for difference in tissue composition between subjects, we performed correction of all metabolite concentrations measured using MEGA-sSPECIAL and sSPECIAL based on the tissue composition in the voxel estimated from brain segmentation results using SPM12 (https://www.fil.ion.ucl.ac.uk/spm/software/spm12/) and transverse (T_2_) relaxation time and tissue-specific water concentration values in [34]. T_2_ relaxation times of all metabolites used for correction can be found in **Supplementary Materials**.

### Functional and structural MRI data processing and analysis

Resting-state fMRI data were processed using a custom pipeline including removal of the first 5 time points, skull-stripping, distortion correction, normalization, smoothing, denoising using ICA-AROMA, white matter and CSF signal regression and high-pass filtering. Detailed pre-processing steps can be found in **Supplementary materials**.

The processed fMRI images were then used as inputs to a standard group-ICA followed by dual-regression procedure to identify canonical resting-state networks (RSNs) at individual level [11]. In this step, the functional data from all subjects and all sessions were first concatenated to create a single 4D dataset. Group-ICA with 25 components were then performed on this dataset and RSNs of interest (i.e. sensorimotor network) were identified using spatial correlations against previously defined maps [35]. We further chose visual network as a control network that is believed to be not as relevant to balance learning to confirm specificity of the learning effect. The number of independent components was chosen in agreement with previous RSFC studies [11], [36]. Next, a dual-regression approach was used to identify the subject-specific RSN maps [37]. This process involved using all 25 components to perform a spatial regression against each separate fMRI dataset and then using the resulting normalized time-course matrices to perform a temporal regression to estimate subject-specific maps that reflect the subject specific strength of functional connectivity. In these subject-specific spatial maps, voxel values represent the strength of association with the particular group ICA of interest. The resulting subject specific RSN map was then masked by the group mean RSN map and the mean value within the networks of interest was extracted for each subject, used as a measure of the strength of functional connectivity within the RSN.

### Diffusion MRI data processing and analysis

DWI data was processed using a custom pipeline including skull-stripping, denoising, Gibbs ringing artifact correction, distortion correction and eddy current correction. Following preprocessing, probabilistic streamline tractography with 10 million streamlines was performed using MRtrix3 [38]. The left and right part of the sensorimotor network defined using group-ICA in the fMRI analysis were used to seed the streamlines, resulting mainly in streamlines passing through the posterior part of corpus callosum and connecting bilateral motor areas. Corpus callosum was used as an inclusion region to avoid spurious streamlines. Sensorimotor network structural connectivity (SNSC) was defined as sum of streamline weights. Detailed processing steps can be found in **Supplementary materials**.

We further calculated whole-brain fractional anisotropy (FA) maps by performing a diffusion tensor imaging (DTI) analysis. FA is a measure of variance of voxel-wise eigenvalues of diffusion directions and is thought to reflect microstructural integrity of white matter, thus providing complementary information to long-range streamline analysis [39] although this metric has been shown to be sensitive but with limited specificity [40]. The resulting FA values were then sampled along the reconstructed white matter pathways connecting the sensorimotor network via MRtrix3’s tck2connectome, yielding mean FA values along the pathway. A demonstration of all MR protocols used in this study is shown in **Figure 1**.

**Figure 1.**
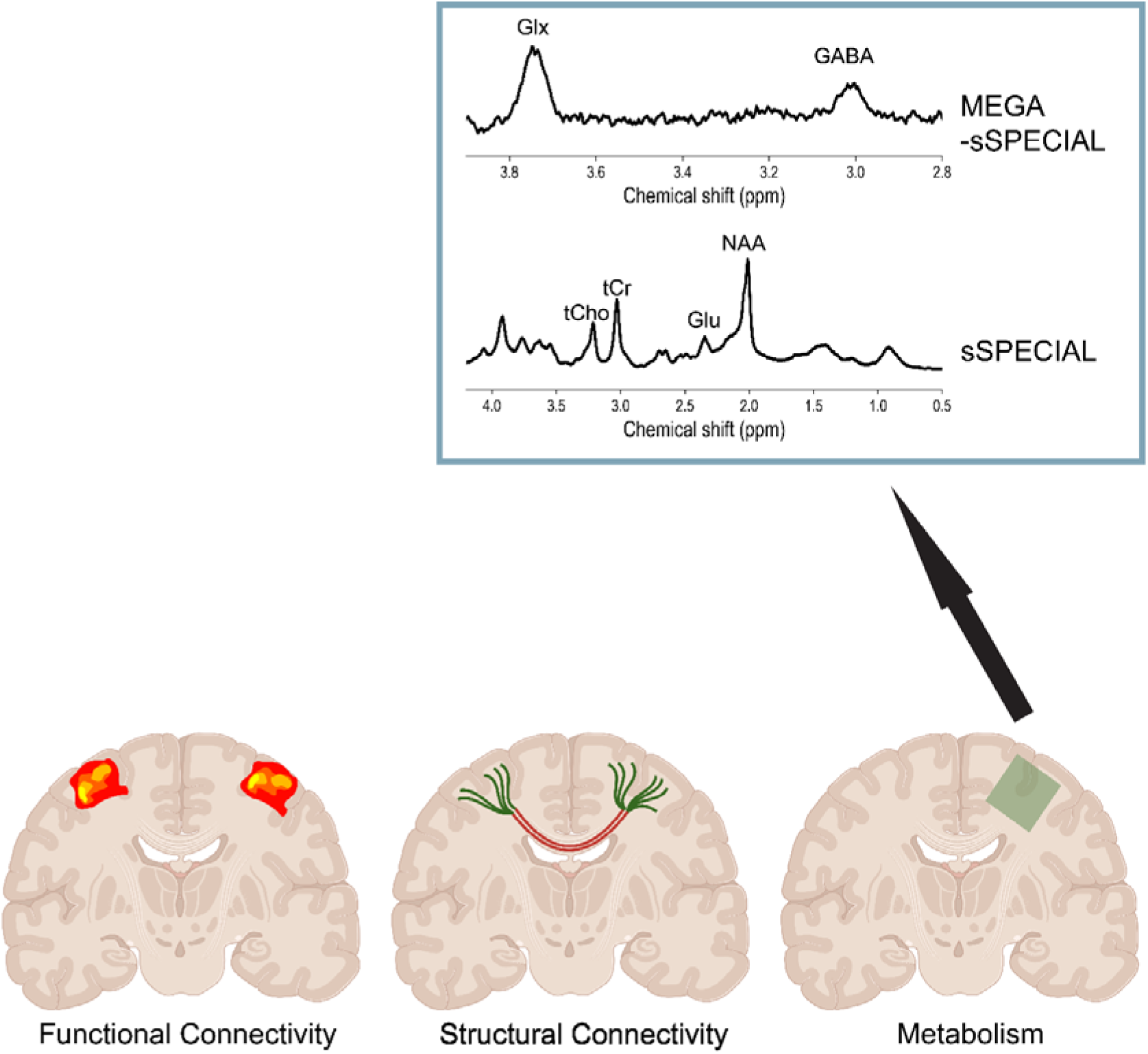
Demonstration of neuroimaging measurement modalities. Left: functional connectivity of SN determined using group-ICA. Mean connectivity strength within the network was extracted for each subject. Middle: structural connectivity of SN determined by probabilistic tractography. The SNSC is determined by multiplying sum of streamline weights by a subject-specific proportionality coefficient determined by SIFT2 algorithm. Right: left sensorimotor cortex metabolism determined by single-voxel MRS. Exemplar spectra of two measurement sequences (MEGA-sSPECIAL and sSPECIAL) are displayed. Figure created with BioRender (https://www.biorender.com/).

### Short interval intracortical inhibition (SICI) measurement

Intracortical inhibition was assessed using a paired-pulse transcranial magnetic stimulation (TMS) protocol. Biphasic suprathreshold single-pulses alternated with suprathreshold pulses preceded by a subthreshold pulse (paired-pulses). The initial pulse of the paired-pulse stimulation served as a conditioning pulse, which activates intracortical inhibitory interneurons. This activation lowers the motor evoked potential (MEP) in the target muscle compared to the unconditioned stimulation [41], [42]. SICI was shown to be a cortical phenomenon [43], [44], [45] relying on GABA_A_ergic mechanisms [46]. SICI measurements were performed while participants balanced on the highest achieved level of the wobble boards during performance measurements at pre-measurement. A figure-of-eight coil (D-B80, Magventure A/S, Farum, Denmark) connected to a MagPro X100 stimulator equipped with the MagOption system (MagVenture A/S, Farum, Denmark) was used. The current was applied in an anterior-posterior direction by orienting the coil handle towards the rear. The optimal location in the left motor cortex to stimulate the right tibialis anterior (TA) muscle was defined in a seated position and marked with color on the scalp. While balancing, the coil was kept in place with a custom-made helmet, which was attached to the ceiling [21]. The active motor threshold (aMT) was determined while standing and was set at the lowest intensity where three out of five consecutive stimulations produced a MEP with a peak-to-peak amplitude equal to or higher than 100 μV [47]. The initial protocol was always a test pulse intensity of 120% of the aMT with a conditioning pulse intensity of 70% of the aMT. If this protocol resulted in inhibition lower than 30% in standing position, the intensities of the pulses were adjusted individually (conditioning pulse: 65% - 80%, test pulse: 120% - 125%). For the post-measurement the same relative intensities of the aMT were employed as for the pre-measurement, however, the aMT was re-determined in the same way as in the pre-test. The interstimulus interval for the paired-pulse stimulations was set at 2 ms [48], [49]. The stimuli were administered in two sets of 20 each, in which single- and paired-pulse stimulations were alternatively applied with an interstimulus pause of 4 seconds. Detailed implementation of SICI measurements can be found in **Supplementary materials**.

### Electromyography (EMG)

After preparing the skin through shaving, abrasion, and disinfection, two surface electrodes (Ag/AgCl, Ambu Blue Sensor N, Ballerup, Denmark) were placed over the muscle belly of the right TA to record bipolar EMG. The EMG electrodes were connected to a transmitter (Myon aktos, Myon AG, Schwarzenberg, Switzerland), amplified analogously (x1000) and wirelessly transmitted to the receiver station of the aktos system. The EMG data were recorded at a sampling frequency of 4 kHz with a custom-made, python-based graphical user interface.

### SICI data processing and analysis

Recorded data were analyzed offline using Matlab. The MEPs were filtered within the range of 20 to 450 Hz and analyzed between 10 to 85 ms after the stimulation. To obtain the single-pulse and paired-pulse MEPs, peak-to-peak amplitudes (mV) were determined. Average SICI values were calculated as the percentage difference between conditioned paired-pulse and unconditioned single-pulse MEPs using the formula 100-(paired-pulse/single-pulse*100) [21], [50]. MEPs were excluded from data analyses if the corresponding background EMG activity before the stimulation was more than 3x the median absolute deviation (MAD) away from the median of all background activities of the corresponding measurement or if the MEPs were smaller than 0.05 mV [51]. Background EMG activity for each MEP was calculated by determining the average root mean square value across a 100 ms interval prior to stimulation [21]. We also performed a control analysis on background EMG in the control and intervention group between pre- and post-tests to ensure that background EMG did not influence the MEP amplitudes.

### Statistical analysis

Statistical analysis was carried out using *statsmodels* package in Python [52]. We tested whether there is a change in outcome variables in the training group compared to the control group using a linear mixed-effect model (LME), which models fixed and random effects simultaneously. In our case, the fixed effects were main effects of group and time as well as their interaction, the random effects are subject-specific random intercepts, and the dependent variable is the measured neuroimaging or performance parameters. In this study, we were mainly interested in the interaction effect, i.e., whether the training changed outcomes compared to control group. Post-hoc two-sample t-tests were followed to determine the direction of change, multiple comparisons were corrected using Benjamini-Hochberg procedure [53]. Cohen’s d was used to report effect sizes. As an explorative analysis, the analysis was also applied to concentrations of other metabolites detected using sSPECIAL to investigate potential effects on other brain metabolites.

In order to test our hypothesis that GABAergic processes (i.e., SICI and GABA-levels) are related to changes in balance performance and functional connectivity, Spearman correlation analyses were performed at baseline and by calculating the changes (delta values) between pre- and post-training values for the four pairs of variables. Correlation analysis was also performed between functional connectivity and balance performance. The Benjamini & Hochberg method (q < 0.05) was used to control the false discovery rate (FDR).

## Results

### Balance performance and SICI

Results of comparing balance performance (area in cm^2^) and SICI between pre and post measurements in both groups are shown in **Figure 2**. LME found an insignificant Group × Time interaction effect when comparing balance performance before and after learning (z_(68)_ = 1.952, p = 0.051, [−0.129, 62.856]). The following t-test revealed that there is a significant decrease in the mean sway area (thus improvement in balance performance) in the learning group (corrected p = 0.0036, t_(17)_ = 3.68, [17.96, 65.93], Cohen’s d = 0.75) and no significant change in the control group (corrected p = 0.36, t_(17)_ = 0.93, [−13.38, 34.54], Cohen’s d = 0.11). Using the time duration between the last training session and measurement as a covariate, we found a significant pre vs. post effect for balance performance (F = 5.79, corrected p = 0.042), indicating significant improvement in performance.

**Figure 2.**
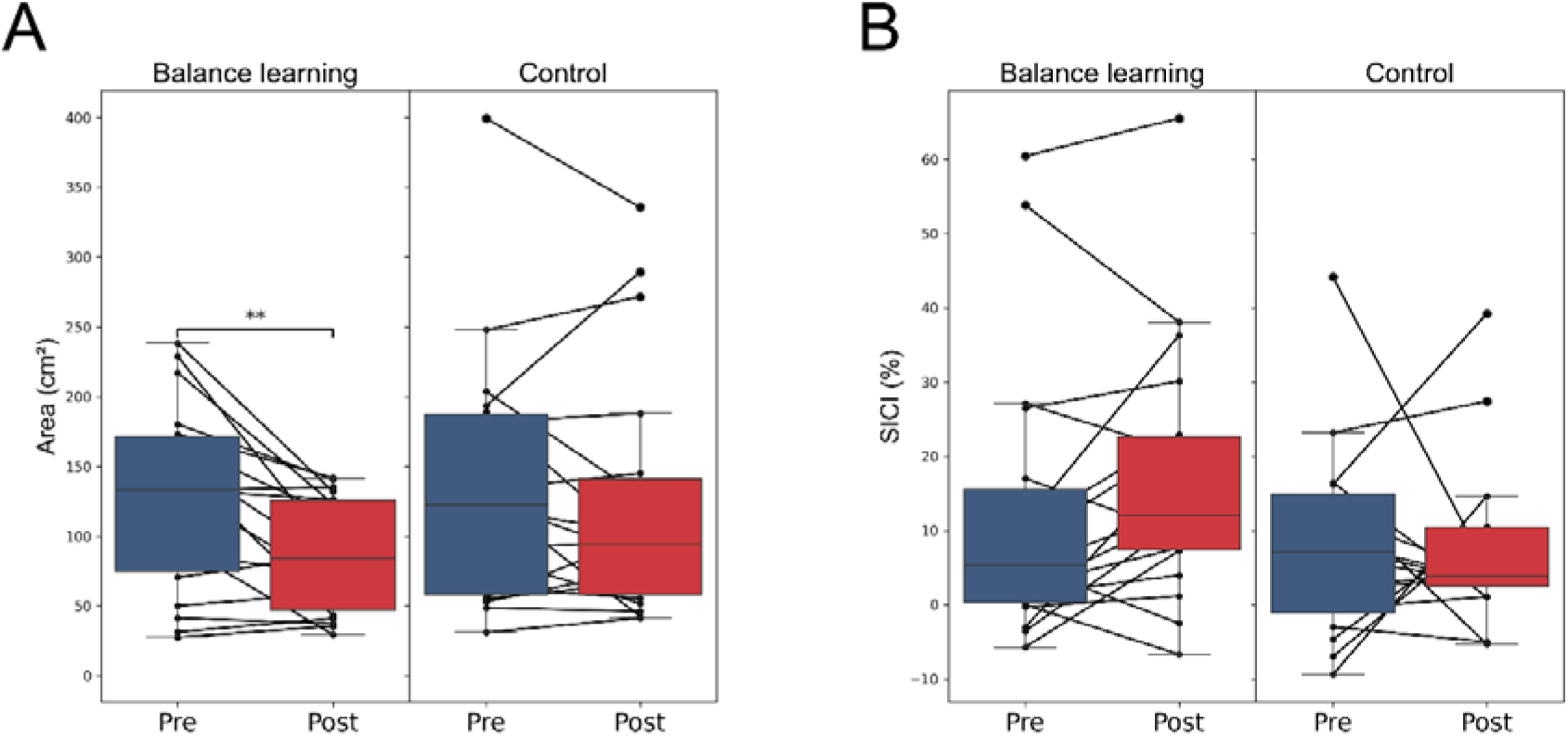
Comparing balance performance (A) and (B) SICI between pre- and post-measurements in balance learning group and control group.

Comparing SICI, we found no significant interaction effect (z_(64)_ = −1.192, p = 0.233, [−14.825 3.610]).

Only 5.8% of the single pulse and 5.9% of the total number of paired-pulse stimulations were excluded from the data analyses to ensure that background activation during the last 100 ms prior to the stimulation did not differ more than three median absolute deviations from the median. There was no significant difference in background activation between the pre- and the post-test in the two groups (both p > 0.05).

### Metabolic plasticity in the sensorimotor cortex

Exemplary voxel placement for MRS measurements is shown in **Figure 3(A)**. LME revealed a significant Group × Time interaction (z_(54)_ = −2.20, p = 0.028, [−0.658, −0.038]) for GABA levels measured by MEGA-sSPECIAL. Furthermore, there was a significant effect of Time (p < 0.001) but no Group effect (p = 0.415). Post-hoc tests confirmed that this effect was driven by increases in GABA levels in the balance learning group (corrected p = 0.006, t_(13)_ = −3.57, [−0.75, −0.18], Cohen’s d = 1.17) (**Figure 3(B)**). There was no difference in GABA levels between pre- and post-measurements for the control group (corrected p = 0.48, t_(13)_ = - 0.72, [−0.26, 0.13], Cohen’s d = 0.13).

**Figure 3.**
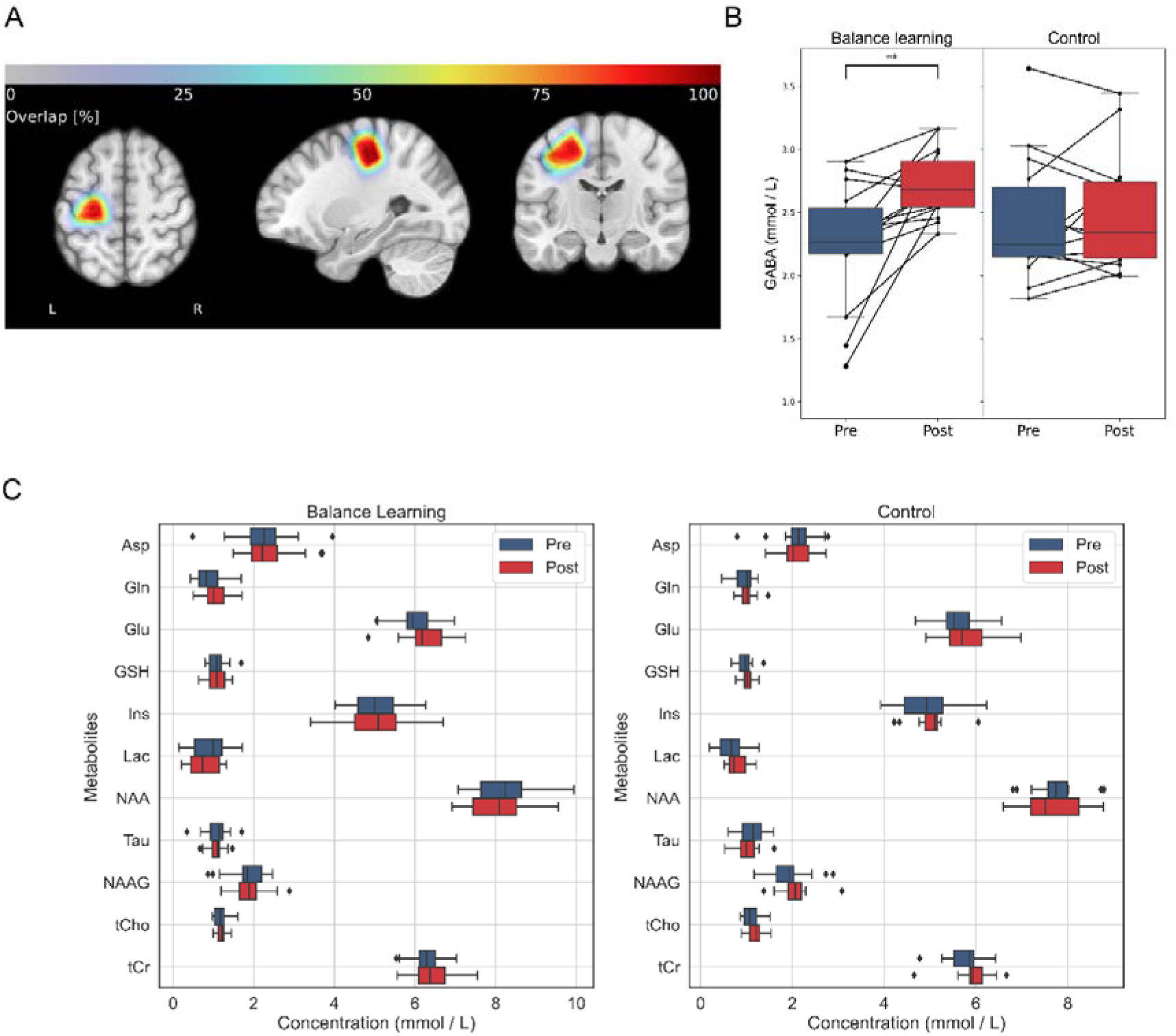
(A) Overlay of voxel placements from all scans for single-voxel MRS created using Osprey [79]. (B) Comparison of sensorimotor GABA levels detected using MEGA-edited MRS between pre and post measurements in balance learning group and control group (**p < 0.01). (C) Comparison of sSPECIAL detected metabolite concentrations before and after training in the two groups. No significant difference is found between the two measurements in any metabolites (uncorrected p > 0.05).

To ensure that our findings are specific to GABA changes, we performed the following control analyses. First, we calculated the signal-to-noise ratio (SNR) using the magnitude of the GABA peak at 3.0 ppm divided by the standard deviation (SD) of the noise signal between 9.5 and 10 ppm. We then used the residual of GABA levels after regressing out SNR as the outcome variable in LME and found similar interaction effects (z_(54)_ = −2.27, p = 0.023, [−0.641, −0.047]). Second, we used the Glx (Glutamate + Glutamine) signal, being another major peak present in MEGA-editing spectra at around 3.8 ppm, as the outcome variable in LME fitting and found no interaction effect (p = 0.49). These control analyses ensured that our findings were specific to GABA levels.

Short echo-time SPECIAL detected 11 main metabolites with mean CRLB < 30% across all groups, which were then included in the explorative analysis. Using all metabolites detected at short echo-time as outcome variable to fit the LME, we found no metabolites showing significant interaction effect (**Figure 3C**). Detailed information about metabolite concentrations measured in this method can be found in **Supplementary Materials**.

### Functional plasticity of the sensorimotor network

In the ICA analysis, two components were identified as belonging to the sensorimotor network, corresponding to ventral and dorsal subdivisions of the network respectively. The ventral sensorimotor network (VSN) is believed to be mainly responsible for motor representations of the upper limb and face, and the dorsal sensorimotor network (DSN) is more closely linked to motor functioning of the lower limb [35]. Both networks were selected as networks of interest for further analysis. One network located mainly in occipital areas was identified as visual network (VN) as a control network. A demonstration of these networks can be found in **Figure 4**.

**Figure 4.**
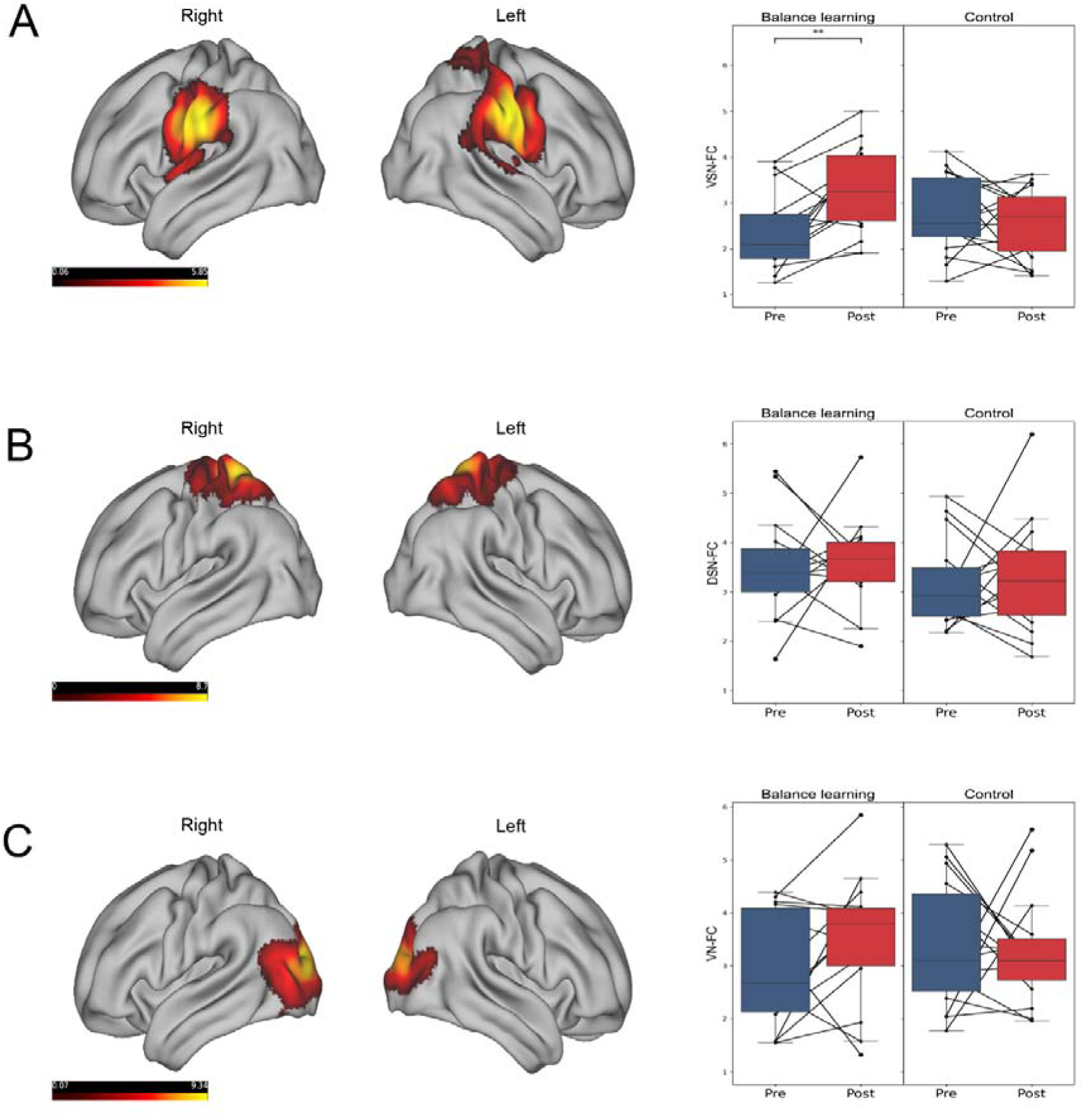
Resting-state functional networks rendered on brain surface using Connectome Workbench (https://www.humanconnectome.org/software/connectome-workbench), detected using group-ICA on the dataset consisting of all scans and all subjects. The values in colormap represent connectivity strength. (A) Left: location of the ventral sensorimotor network. Right: comparison of functional connectivity (FC) between groups, we found a significant improvement in VSN-FC in balance learning group but not in control group (**p < 0.01). (B) Location of the dorsal sensorimotor network (DSN) and between-group comparisons. (C) Location of the visual network (VN) as a control network and between-group comparisons. We did not find significant between-group differences for DSN-FC and VN-FC.

In the next step, balance learning induced changes in the SNFC were tested. Taking the mean connectivity strength within the network as a summary metric, for ventral sensorimotor network functional connectivity (VSN-FC), we found significant Group × Time interaction effect in LME fitting (z_(61)_ = −3.11, p = 0.002, [−1.871, −0.424]) and a significant main effect for Time (p < 0.001) but no Group effect (p = 0.245). Post-hoc t-tests further confirmed that this is driven by a significant increase in VSN-FC in the balance learning group (corrected p = 0.006, t_(13)_ = −3.63, [−1.54, −0.39], Cohen’s d = 1.07) while no change occurred in the control group (corrected p = 0.37, t_(15)_ = 0.91, [−0.32, 0.8], Cohen’s d = 0.29; **Figure 4A**). We did not find a significant interaction effect for dorsal sensorimotor network functional connectivity (DSN-FC) (z_(61)_ = −0.183, p = 0.855, [−1.039, 0.862]). As a control analysis, we did not find a significant interaction effect for visual network functional connectivity (VN-FC) neither (z_(61)_ = −1.071, p = 0.284, [−1.620, 0.475]) (**Figure 4B, C**).

### Structural plasticity of the sensorimotor network

We used probabilistic tractography to test whether long-term balance learning induced alterations in the brain’s structural connectivity. White matter bundles between bilateral VSN were retrieved and structural connectivity (SC) was defined as a weighted sum of streamline counts. Using this summary metric as the outcome variable in the LME, we found no significant effects for Group × Time (z_(58)_ = −1.486, p = 0.137, [−0.073, 0.010]) (Supplementary Figure 2A).

In order to assess potential microstructural alterations induced by learning, we extracted mean FA values along the tracts connecting bilateral motor areas. LME analysis using FA values as outcome variables did not reveal significant interaction effect of Group × Time (z_(58)_ = 1.026, p = 0.305, [−0.006, 0.019]) (Supplementary Figure 2B).

### Correlation analysis

Since only VSN-FC showed between group differences induced by learning, we hereby focused on VSN-FC for the following correlation analysis. Correlation analysis between SICI/GABA and VSN-FC/balance performance are shown in **Figure 5**. At baseline, there was a significant association between SICI and VSN-FC (uncorrected p = 0.01, r = 0.48). In addition, we found a significant association between higher VSN-FC and better balance performance (uncorrected p = 0.04, r = −0.37) at baseline. In the learning group, the improvement in balance performance indicated by the reduced mean sway area was significantly correlated with upregulation of SICI (uncorrected p = 0.02, r = −0.54). However, none of these correlations survived the FDR correction (corrected p > 0.05).

**Figure 5.**
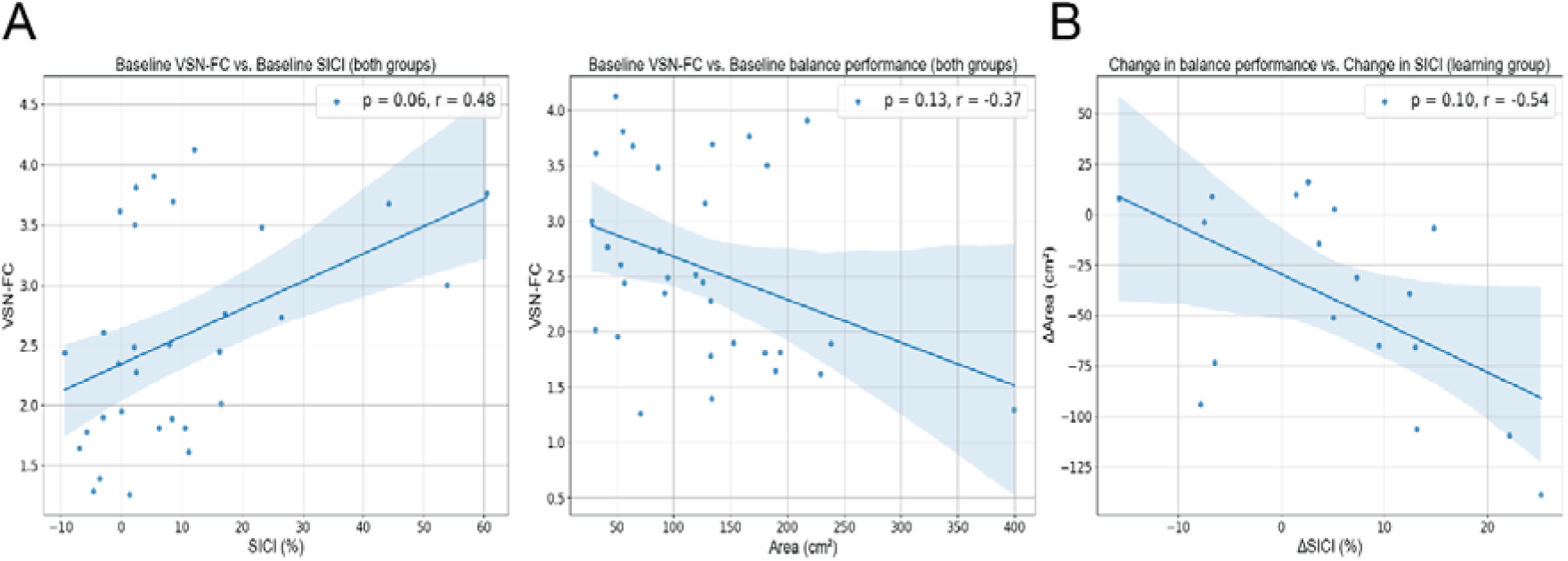
Correlations between the level of intracortical inhibition/GABA on one hand and VSN-FC and balance performance on the other hand were investigated both for baseline values in both groups (A), for delta values in learning group (B). The three correlations were significant but did not survive FDR correction. All p-values are presented as corrected. Other correlations involving GABA were not significant before correction and the plots can be found in supplementary Figure 4.

## Discussion

In this study, we employed multi-model neuroimaging and neurophysiological measurements to investigate the modulatory effects of long-term balance learning on cortical inhibition, functional and structural connectivity among an elderly cohort. After three months of balance learning, subjects significantly upregulated sensorimotor cortical GABA levels and VSN-FC compared to a passive control group. These findings shed light on the adaptation of cortical inhibitory processes induced by motor learning and their interplay with behavior and brain plasticity.

### Upregulation of sensorimotor cortical GABA and functional connectivity

Three months of balance learning significantly improved sensorimotor GABA level in the balance learning group compared to the control group. In previous studies that examined motor learning induced plasticity of the GABAergic system, it has often been reported that MRS-measured GABA levels in sensorimotor cortex would be reduced over the learning period [54], [55], suggesting that reduction in cortical inhibitory tone is instrumental in promoting plasticity processes in motor system. As our findings seem to be contradictory at first sight, it is important to highlight that decreases in GABA reported in these previous studies were mainly observed during the very early stage of motor learning (i.e., on the same day). During the initial acquisition phase, a decrease in GABA is not surprising as it is known that reduction in GABA inhibition facilitates long-term potentiation (LTP)-like activity in the motor cortex thereby “unmasking” existing horizontal connections which allows rapid changes in sensory or motor representations [54]. In our longitudinal setting, however, the initial plasticity processes needing reductions in GABA have most likely already been completed. Thus, the observed increases in GABA after 3 months of training may actually represent the more relevant and outlasting change within the cortical inhibitory system.

Another important factor to consider when interpreting the observed increase in GABA is that in contrast to most previous studies, our study population consisted of elderly adults. Previous *in vivo* studies reported that older adults experience aging-related cortical neural de-differentiation, and the extent of the decline in neural distinctiveness correlates with reductions in cortical GABA levels [56], [57]. That is, subjects with lower GABA levels show less distinctive neural activation patterns in response to external stimuli. Thus, the increased GABA found in the current study may reflect long-term learning-induced improvements in neural differentiation and selectivity. This explanation aligns with a previous study linking higher GABA levels to better motor performance, specifically in older adults [24]. Furthermore, the absence of changes in other metabolites such as NAA and Cr suggest that balance learning mainly involves plasticity at the synaptic and network levels, rather than substantial alterations in the overall neuronal density or energy metabolism.

In line with previous studies [22], [58], [59], we found that three months of balance learning improved functional connectivity within the sensorimotor network. A decline in within-network connectivity compared to inter-network connectivity has been found with aging [7], suggesting that aging leads to reduced specificity of sensorimotor brain functions. Reduced SNFC has also been linked to motor deficits [4] and declined levels of neural distinctiveness [60]. Therefore, the observed changes in SN functional connectivity, in conjunction with the increase in GABA levels associated with neural distinctiveness, support the hypothesis that balance learning alleviates aging-related neural de-differentiation. In addition, deficits in SNFC have been found in a series of neurophysiological diseases, such as motor dysfunction in stroke [61], bipolar disorder [62] and Alzheimer’s disease [63]. Thus, our findings may indicate the potential relevance of balance learning as a therapeutic strategy for elderly adults and people suffering from neurological diseases that involve impairments in motor inhibition and functional connectivity. Interestingly though, we found that the observed changes in SNFC induced by balance learning were located in the upper limb, face and trunk areas of the motor cortex (VSN-FC) but not in areas representing the lower limb (DSN-FC). Also, the VSN-FC at baseline correlated with balance performance. This latter finding echoes several previous studies: Taube et al. reported brain activation within the primary motor cortex only in these ventral motor areas during motor imagery and action observation of balance tasks [30], and Taubert et al. reported increased ventral premotor cortex functional connectivity over the course of six weeks of balance learning [16]. Although it may seem intuitive that balance learning involves relatively more leg areas, current results suggest that balance may also be represented in the ventral part of motor cortex. One potential explanation is that balancing involves not only leg movements but whole-body coordination. For example, Objero et al. reported that restricting arm movements could greatly influence balance performance [64]. Thus, future study is still needed to investigate the representation of balancing in the motor cortex.

In this study, we restricted our analysis of white matter connectivity and microstructure to the posterior part of corpus callosum connecting bilateral motor areas, to keep in line with functional connectivity analysis. Interestingly, we did not observe any significant changes in structural connectivity or white matter microstructure within the region. The absence of changes echoes a previous review suggesting that while corpus callosum may play a supportive role in balance, it likely is not the most critical underlying component of balance in the brain, as there is a lack of findings of balance learning induced changes in this area [65]. Possible extensions include tract-based analyses across the entire WM, including the bilateral cortico-spinal tracts, as well as the investigation of further diffusion metrics beyond FA, such as DKI scalars [66] or parameters of the WM two-compartment model [67], [68], [69]. Future study is therefore needed to investigate structural underpinnings of GABA and functional connectivity changes in balance learning.

### Adaptations in cortical inhibition induced by balance learning

In previous studies, upregulation in SICI has been observed after motor learning. For example, Cirillo et al. [70] showed upregulation in SICI after sequential visual isometric training, indicating that enhanced GABA-mediated inhibition may promote consolidation of the newly acquired skill. Mouthon et al. [21] showed in young healthy adults that SICI was increased after learning a balance task for two weeks, and the upregulation in SICI was correlated with improved performance. Similar to these findings, our data points to a positive association between changes in SICI after long-term balance learning and improvements in balance performance, but among an elderly cohort. Thus, complementary to these studies, our results provide a fresh perspective that upregulation of cortical inhibition can also be induced in elderly adults, a population known to have cortical inhibition deficits [71], [72]. Our results suggest that balance learning has the potential to counteract aging-related decline in cortical inhibition and its associated motor function deficits.

Interestingly, MRS-derived GABA levels in the sensorimotor cortex were augmented consistently in the trained subjects but no correlation to balance performance or functional connectivity was apparent. It is worth noting that MRS measures the local GABA concentration within a specific region without distinguishing between cytoplasmic, vesicular and extra-synaptic GABA pools [26]. As mentioned earlier, MRS-assessed GABA may reflect learning-induced improvements in neural differentiation and selectivity. On the contrary, SICI specifically detects the activation of GABA_A_ receptors during balancing (as compared to the resting position in the scanner) and can thus more accurately quantify the degree of cortical inhibition, which involves GABA acting on synaptic GABA_A_ receptors [26] during task execution. Thus, we propose that measures of SICI and MRS are two complementary methods to assess GABAergic mechanisms. This is based on the assumption that MRS likely measures tonic inhibition driven by extracellular GABA acting on extrasynaptic GABA_A_ receptors whereas SICI depicts vesicular GABA acting on synaptic GABA_A_ (for review see [73]). Accordingly, we propose the idea that MRS-derived GABA-levels in the primary motor cortex indicate the more general ‘capacity for inhibition’ whereas assessment of SICI during motor execution reveals more specifically the ‘ability to modulate inhibition’ for a specific task and muscle. Both the ‘capacity of inhibition’ and the ‘ability to modulate inhibition’ can be changed through training as indicated by the present results. However, the global increase in the level of total GABA (MRS-measure) can be assessed at rest, while the ability to modulate inhibition becomes only apparent by TMS-measurements during execution of the trained motor task (i.e., SICI measures during balancing). This interpretation aligns with previous studies which reported that adaptation in SICI were not seen when assessed in a control muscle or in the trained muscle at rest [20], [74], [75].

### Associations between cortical inhibition, functional connectivity and behavior

The correlation analyses suggests a positive association between VSN-FC and intracortical inhibition, which echoes a previous study reporting that during the earliest stages of sensorimotor learning, GABA levels are linked on a subject-level basis to both, behavioral learning and a strengthening of functional connections that persists beyond the training period [8]. In another study [10], GABA levels within the regions in the default mode network (DMN) and control network were found to be significantly correlated with de-activation of the DMN and DMN-CN correlation. This suggests that GABAergic processes are involved in orchestrating between-network brain activity at rest and during task performance. Taken together, the findings are consistent with the idea that GABA-mediated inhibition is linked to maintenance of newly learned information and further validates the view that GABAergic processes may play an important role in modulating functional coupling between brain regions. However, the correlation did not survive FDR correction, thus our interpretation is needed to be validated in a larger sample size.

### Limitations

One potential methodological concern of the current study is that the TMS and MRS measurement did not cover exactly the same spot in the primary motor cortex. However, we consider this not as a limitation as we did not expect to see “singular adaptations only for the tibialis anterior muscle” in response to learning balance tasks. We have chosen the tibialis muscle because of its high degree of cortical control [74], [76], susceptibility to age-related declines [71], importance for balance control [77], good accessibility by TMS [74], [76] and very precise and task-specific modulation of intracortical inhibition [74]. From a biomechanical point of view, other (antigravity) muscles might be more important to ensure upright stance such as the soleus muscle but the latter muscle is less well cortically controlled and exhibits less task-specific modulation in SICI in general and with age in particular. Furthermore, the entire leg and trunk areas as well as the arms are supposed to strongly contribute when learning balance but are less well accessible and reliably when measured with TMS. The tibialis anterior muscle represents a reliable candidate within the primary motor cortex to demonstrate the effect of balance training by means of TMS (SICI) because it modulates intracortical inhibition very well. In contrast, the previously identified spot in the ventral motor cortex [30] using fMRI seemed to be most promising for assessing more global changes in the primary motor cortex.

Another potential limitation in our study is that the control group did not undergo any control intervention. Thus, the observed pre-post changes could have resulted from other effects such as motivation and additional social contact than the balance training per se. However, we believe that the changes seen after the balance training are due to the balance training itself. This is because the observed changes in SICI are in line with previous research in younger [20], [21] as well as older subjects (Kuhn et al. 2024, under revision). With respect to motivation, i.e., subjects wanted to perform better in the post-than the pre-test, this would not have affected the resting MRS and MRI as well as the SICI measurements. Given the duration of the training and thus, the time elapsed between the pre- and the post-test, it is very unlikely that retest effects played an important role. This is supported by the non-significant changes in the control group. It is true that the balance training was organized in group sessions which likely has led to additional social contact, but this probably only increased the motivation to participate in the training without effecting the pre-vs post-test performance.

In addition, the current study is not pre-registered, which may raise doubts about the analysis plan. However, we conducted a prior sample size calculation based on the SICI and performance improvement from our just published results [78]. The detailed calculation is provided in supplementary materials sample size calculation section. Future research should consider pre-registering their studies before launching to mitigate such concerns.

## Conclusion

In summary, the present study employed a multi-modal neuroimaging and TMS approach to systematically investigate the effects of balance learning on the cortical sensorimotor system of older adults under a longitudinal framework. We found that three months of balance learning led to increased GABA levels in the sensorimotor cortex and enhanced functional connectivity of the sensorimotor network. Furthermore, correlation analysis suggests a potential positive association between baseline VSN-FC and balance performance, between baseline VSN-FC and SICI, and between improvements in balance performance and upregulation in SICI, though these correlations did not survive the FDR correction. This study provides novel insights into the neural plasticity associated with the GABAergic system in elderly adults and suggests that balance learning has the potential to counteract aging-related motor and functional connectivity decline. We suggest that changes in the amount of GABA in the sensorimotor cortex (i.e., measured with MRS) indicate the ‘capacity for inhibition’ (the resources for inhibition) whereas changes in intracortical inhibition (SICI) highlights the ‘ability to modulate inhibition’ and the ‘ability to shape functional connections across brain regions’ (functional connectivity).

## Acknowledgements

This work was supported by the Swiss National Science Foundation (grants n° 32003B_197687). We acknowledge access to the facilities and expertise of the CIBM Center for Biomedical Imaging, a Swiss research center of excellence funded and supported by Lausanne University Hospital (CHUV), University of Lausanne (UNIL), Ecole Polytechnique Fédérale de Lausanne (EPFL), University of Geneva (UNIGE) and Geneva University Hospitals (HUG).

## Data availability

All data used in this study is available in **Supplementary Data**.

## References

[1] R. D. Seidler et al., “Motor control and aging: Links to age-related brain structural, functional, and biochemical effects,” Neurosci. Biobehav. Rev., vol. 34, no. 5, pp. 721–733, 2010, doi: 10.1016/j.neubiorev.2009.10.005.

[2] O. Levin, H. Fujiyama, M. P. Boisgontier, S. P. Swinnen, and J. J. Summers, “Aging and motor inhibition: A converging perspective provided by brain stimulation and imaging approaches,” Neurosci. Biobehav. Rev., vol. 43, pp. 100–117, 2014, doi: 10.1016/j.neubiorev.2014.04.001.

[3] L. Hermans et al., “Brain GABA levels are associated with inhibitory control deficits in older adults,” J. Neurosci., vol. 38, no. 36, pp. 7844–7851, 2018, doi: 10.1523/JNEUROSCI.0760-18.2018.

[4] K. Cassady et al., “Sensorimotor network segregation declines with age and is linked to GABA and to sensorimotor performance,” Neuroimage, vol. 186, no. November 2018, pp. 234–244, 2019, doi: 10.1016/j.neuroimage.2018.11.008.

[5] B. Calvo-Flores Guzmán, C. Vinnakota, K. Govindpani, H. J. Waldvogel, R. L. M. Faull, and A. Kwakowsky, “The GABAergic system as a therapeutic target for Alzheimer’s disease,” J. Neurochem., vol. 146, no. 6, pp. 649–669, 2018, doi: 10.1111/jnc.14345.

[6] M. J. Firbank et al., “Reduced occipital GABA in Parkinson disease with visual hallucinations,” Neurology, vol. 91, no. 7, pp. e675–e685, 2018, doi: 10.1212/WNL.0000000000006007.

[7] M. Y. Chan, D. C. Park, N. K. Savalia, S. E. Petersen, and G. S. Wig, “Decreased segregation of brain systems across the healthy adult lifespan,” Proc. Natl. Acad. Sci. U. S. A., vol. 111, no. 46, pp. E4997–E5006, 2014, doi: 10.1073/pnas.1415122111.

[8] F. T. van Vugt, J. Near, T. Hennessy, J. Doyon, and D. J. Ostry, “Early stages of sensorimotor map acquisition: Neurochemical signature in primary motor cortex and its relation to functional connectivity,” J. Neurophysiol., vol. 124, no. 6, pp. 1615–1624, 2020, doi: 10.1152/jn.00285.2020.

[9] N. Levar, T. J. Van Doesum, D. Denys, and G. A. Van Wingen, “Anterior cingulate GABA and glutamate concentrations are associated with resting-state network connectivity,” Sci. Rep., vol. 9, no. 1, pp. 1–8, 2019, doi: 10.1038/s41598-018-38078-1.

[10] X. Chen et al., “Regional GABA Concentrations Modulate Inter-network Resting-state Functional Connectivity,” Cereb. Cortex, vol. 29, no. 4, pp. 1607–1618, 2019, doi: 10.1093/cercor/bhy059.

[11] C. J. Stagg et al., “Local GABA concentration is related to network-level resting functional connectivity,” Elife, vol. 2014, no. 3, pp. 1–9, 2014, doi: 10.7554/eLife.01465.

[12] C. Rosano, H. Aizenstein, J. Brach, A. Longenberger, S. Studenski, and A. B. Newman, “Gait measures indicate underlying focal gray matter atrophy in the brain of older adults,” Journals Gerontol. - Ser. A Biol. Sci. Med. Sci., vol. 63, no. 12, pp. 1380–1388, 2008, doi: 10.1093/gerona/63.12.1380.

[13] B. Draganski, C. Gaser, V. Busch, G. Schuierer, U. Bogdahn, and A. May, “Changes in grey matter induced by training,” Nature, vol. 427, pp. 311–312, 2004, [Online]. Available: www.nature.com/nature

[14] C. D. Rowley et al., “Exercise and microstructural changes in the motor cortex of older adults,” Eur. J. Neurosci., vol. 51, no. 7, pp. 1711–1722, 2020, doi: 10.1111/ejn.14585.

[15] L. Bezzola, S. Mérillat, C. Gaser, and L. Jäncke, “Training-induced neural plasticity in golf novices,” J. Neurosci., vol. 31, no. 35, pp. 12444–12448, 2011, doi: 10.1523/JNEUROSCI.1996-11.2011.

[16] M. Taubert, G. Lohmann, D. S. Margulies, A. Villringer, and P. Ragert, “Long-term effects of motor training on resting-state networks and underlying brain structure,” Neuroimage, vol. 57, no. 4, pp. 1492–1498, 2011, doi: 10.1016/j.neuroimage.2011.05.078.

[17] B. Sehm et al., “Structural brain plasticity in parkinson’s disease induced by balance training,” Neurobiol. Aging, vol. 35, no. 1, pp. 232–239, 2014, doi: 10.1016/j.neurobiolaging.2013.06.021.

[18] K. Ueta, N. Mizuguchi, T. Sugiyama, T. Isaka, and S. Otomo, “The Motor Engram of Functional Connectivity Generated by Acute Whole-Body Dynamic Balance Training,” Med. Sci. Sports Exerc., vol. 54, no. 4, pp. 598–608, 2022, doi: 10.1249/MSS.0000000000002829.

[19] S. Magon et al., “Striatal functional connectivity changes following specific balance training in elderly people: MRI results of a randomized controlled pilot study,” Gait Posture, vol. 49, pp. 334–339, 2016, doi: 10.1016/j.gaitpost.2016.07.016.

[20] W. Taube, A. Gollhofer, and B. Lauber, “Training-, muscle- and task-specific up- and downregulation of cortical inhibitory processes,” Eur. J. Neurosci., vol. 51, no. 6, pp. 1428–1440, 2020, doi: 10.1111/ejn.14538.

[21] A. Mouthon and W. Taube, “Intracortical Inhibition Increases during Postural Task Execution in Response to Balance Training,” Neuroscience, vol. 401, pp. 35–42, 2019, doi: 10.1016/j.neuroscience.2019.01.007.

[22] C. Sampaio-Baptista, N. Filippini, C. J. Stagg, J. Near, J. Scholz, and H. Johansen-Berg, “Changes in functional connectivity and GABA levels with long-term motor learning,” Neuroimage, vol. 106, pp. 15–20, 2015, doi: 10.1016/j.neuroimage.2014.11.032.

[23] S. Chalavi et al., “The neurochemical basis of the contextual interference effect,” Neurobiol. Aging, vol. 66, pp. 85–96, 2018, doi: 10.1016/j.neurobiolaging.2018.02.014.

[24] C. Maes, K. Cuypers, K. F. Heise, R. A. E. Edden, J. Gooijers, and S. P. Swinnen, “GABA levels are differentially associated with bimanual motor performance in older as compared to young adults,” Neuroimage, vol. 231, no. January, p. 117871, 2021, doi: 10.1016/j.neuroimage.2021.117871.

[25] S. I. Lim and L. Xin, “γ-aminobutyric acid measurement in the human brain at 7 T: Short echo-time or Mescher–Garwood editing,” NMR Biomed., no. June 2021, pp. 1–17, 2022, doi: 10.1002/nbm.4706.

[26] C. J. Stagg, V. Bachtiar, and H. Johansen-Berg, “What are we measuring with GABA Magnetic Resonance Spectroscopy?,” Commun. Integr. Biol., vol. 4, no. 5, pp. 573–575, 2011, doi: 10.4161/cib.16213.

[27] M. C. Ferland et al., “Transcranial Magnetic Stimulation and H1-Magnetic Resonance Spectroscopy Measures of Excitation and Inhibition Following Lorazepam Administration,” Neuroscience, vol. 452, pp. 235–246, 2021, doi: 10.1016/j.neuroscience.2020.11.011.

[28] J. P. Marques, T. Kober, G. Krueger, W. van der Zwaag, P. F. Van de Moortele, and R. Gruetter, “MP2RAGE, a self bias-field corrected sequence for improved segmentation and T1-mapping at high field,” Neuroimage, vol. 49, no. 2, pp. 1271–1281, 2010, doi: 10.1016/j.neuroimage.2009.10.002.

[29] W. M. Teeuwisse, W. M. Brink, and A. G. Webb, “Quantitative assessment of the effects of high-permittivity pads in 7 Tesla MRI of the brain,” Magn. Reson. Med., vol. 67, no. 5, pp. 1285–1293, 2012, doi: 10.1002/mrm.23108.

[30] W. Taube, M. Mouthon, C. Leukel, H. M. Hoogewoud, J. M. Annoni, and M. Keller, “Brain activity during observation and motor imagery of different balance tasks: An fMRI study,” Cortex, vol. 64, pp. 102–114, 2015, doi: 10.1016/j.cortex.2014.09.022.

[31] S. Moeller et al., “Multiband multislice GE-EPI at 7 tesla, with 16-fold acceleration using partial parallel imaging with application to high spatial and temporal whole-brain FMRI,” Magn. Reson. Med., vol. 63, no. 5, pp. 1144–1153, 2010, doi: 10.1002/mrm.22361.

[32] R. Simpson, G. A. Devenyi, P. Jezzard, T. J. Hennessy, and J. Near, “Advanced processing and simulation of MRS data using the FID appliance (FID-A)—An open source, MATLAB-based toolkit,” Magn. Reson. Med., vol. 77, no. 1, pp. 23–33, 2017, doi: 10.1002/mrm.26091.

[33] S. W. Provencher, “Automatic quantitation of localized in vivo 1H spectra with LCModel,” NMR Biomed., vol. 14, no. 4, pp. 260–264, 2001, doi: 10.1002/nbm.698.

[34] E. Dhamala, I. Abdelkefi, M. Nguyen, T. J. Hennessy, H. Nadeau, and J. Near, “Validation of in vivo MRS measures of metabolite concentrations in the human brain,” NMR Biomed., vol. 32, no. 3, pp. 1–15, 2019, doi: 10.1002/nbm.4058.

[35] B. T. Thomas Yeo et al., “The organization of the human cerebral cortex estimated by intrinsic functional connectivity,” J. Neurophysiol., vol. 106, no. 3, pp. 1125–1165, 2011, doi: 10.1152/jn.00338.2011.

[36] D. V. Smith et al., “Characterizing individual differences in functional connectivity using dual-regression and seed-based approaches,” Neuroimage, vol. 95, pp. 1–12, 2014, doi: 10.1016/j.neuroimage.2014.03.042.

[37] L. D. Nickerson, S. M. Smith, D. Öngür, and C. F. Beckmann, “Using dual regression to investigate network shape and amplitude in functional connectivity analyses,” Front. Neurosci., vol. 11, no. MAR, pp. 1–18, 2017, doi: 10.3389/fnins.2017.00115.

[38] J. D. Tournier et al., “MRtrix3: A fast, flexible and open software framework for medical image processing and visualisation,” Neuroimage, vol. 202, no. January, p. 116137, 2019, doi: 10.1016/j.neuroimage.2019.116137.

[39] A. L. Alexander et al., “Characterization of Cerebral White Matter Properties Using Quantitative Magnetic Resonance Imaging Stains,” Brain Connect., vol. 1, no. 6, pp. 423–446, 2011, doi: 10.1089/brain.2011.0071.

[40] I. O. Jelescu and M. D. Budde, “Design and validation of diffusion MRI models of white matter,” Front. Phys., vol. 5, no. NOV, 2017, doi: 10.3389/fphy.2017.00061.

[41] T. Kujirai et al., “Corticocortical inhibition in human motor cortex,” J. Physiol., vol. 471, no. 1, pp. 501–519, 1993, doi: 10.1016/0006-2952(95)00035-X.

[42] J. C. Rothwell, B. L. Day, P. D. Thompson, and T. Kujirai, “Short latency intracortical inhibition: One of the most popular tools in human motor neurophysiology,” J. Physiol., vol. 587, no. 1, pp. 11–12, 2009, doi: 10.1113/jphysiol.2008.162461.

[43] V. Di Lazzaro and J. C. Rothwell, “Corticospinal activity evoked and modulated by non-invasive stimulation of the intact human motor cortex,” J. Physiol., vol. 592, no. 19, pp. 4115–4128, 2014, doi: 10.1113/jphysiol.2014.274316.

[44] V. Di Lazzaro et al., “Magnetic transcranial stimulation at intensities below active motor threshold activates intracortical inhibitory circuits,” Exp. Brain Res., vol. 119, no. 2, pp. 265–268, 1998, doi: 10.1007/s002210050341.

[45] D. Weise et al., “Microcircuit mechanisms involved in paired associative stimulation-induced depression of corticospinal excitability,” J. Physiol., vol. 591, no. 19, pp. 4903–4920, 2013, doi: 10.1113/jphysiol.2013.253989.

[46] U. Ziemann, J. C. Rothwell, and M. C. Ridding, “Interaction between intracortical inhibition and facilitation in human motor cortex,” J. Physiol., vol. 496, no. 3, pp. 873–881, 1996, doi: 10.1113/jphysiol.1996.sp021734.

[47] P. M. Rossini et al., “Non-invasive electrical and magnetic stimulation of the brain, spinal cord, roots and peripheral nerves: Basic principles and procedures for routine clinical and research application: An updated report from an I.F.C.N. Committee,” Clin. Neurophysiol., vol. 126, no. 6, pp. 1071–1107, 2015, doi: 10.1016/j.clinph.2015.02.001.

[48] S. Vucic, B. C. Cheah, A. V. Krishnan, D. Burke, and M. C. Kiernan, “The effects of alterations in conditioning stimulus intensity on short interval intracortical inhibition,” Brain Res., vol. 1273, pp. 39–47, 2009, doi: 10.1016/j.brainres.2009.03.043.

[49] L. Roshan, G. O. Paradiso, and R. Chen, “Two phases of short-interval intracortical inhibition,” Exp. Brain Res., vol. 151, no. 3, pp. 330–337, 2003, doi: 10.1007/s00221-003-1502-9.

[50] Y. A. Kuhn, M. Keller, J. Ruffieux, and W. Taube, “Adopting an external focus of attention alters intracortical inhibition within the primary motor cortex,” Acta Physiol., vol. 220, no. 2, pp. 289–299, 2017, doi: 10.1111/apha.12807.

[51] M. Keller, W. Taube, and B. Lauber, “Task-dependent activation of distinct fast and slow(er) motor pathways during motor imagery,” Brain Stimul., vol. 11, no. 4, pp. 782–788, 2018, doi: 10.1016/j.brs.2018.02.010.

[52] S. Seabold and J. Perktold, “Statsmodels: Econometric and Statistical Modeling with Python,” Proc. 9th Python Sci. Conf., no. Scipy, pp. 92–96, 2010, doi: 10.25080/majora-92bf1922-011.

[53] Y. Benjamini and Y. Hochberg, “Controlling the False Discovery Rate: A Practical and Powerful Approach to Multiple Testing,” J. R. Stat. Soc., vol. 57, no. 1, pp. 289–300, 1995, doi: 10.2307/2346101.

[54] A. Floyer-Lea, M. Wylezinska, T. Kincses, and P. M. Matthews, “Rapid modulation of GABA concentration in human sensorimotor cortex during motor learning,” J. Neurophysiol., vol. 95, no. 3, pp. 1639–1644, 2006, doi: 10.1152/jn.00346.2005.

[55] J. Kolasinski, E. L. Hinson, A. P. Divanbeighi Zand, A. Rizov, U. E. Emir, and C. J. Stagg, “The dynamics of cortical GABA in human motor learning,” J. Physiol., vol. 597, no. 1, pp. 271–282, 2019, doi: 10.1113/JP276626.

[56] P. Lalwani et al., “Neural distinctiveness declines with age in auditory cortex and is associated with auditory GABA levels,” Neuroimage, vol. 201, no. July, p. 116033, 2019, doi: 10.1016/j.neuroimage.2019.116033.

[57] A. G. Leventhal, Y. Wang, M. Pu, Y. Zhou, and Y. Ma, “GABA and its agonists improved visual cortical function in senescent monkeys,” Science (80-. )., vol. 300, no. 5620, pp. 812–815, 2003, doi: 10.1126/science.1082874.

[58] S. Vahdat, M. Darainy, T. E. Milner, and D. J. Ostry, “Functionally specific changes in resting-state sensorimotor networks after motor learning,” J. Neurosci., vol. 31, no. 47, pp. 16907–16915, 2011, doi: 10.1523/JNEUROSCI.2737-11.2011.

[59] T. Marins, E. C. Rodrigues, T. Bortolini, B. Melo, J. Moll, and F. Tovar-Moll, “Structural and functional connectivity changes in response to short-term neurofeedback training with motor imagery,” Neuroimage, vol. 194, no. July 2018, pp. 283–290, 2019, doi: 10.1016/j.neuroimage.2019.03.027.

[60] K. Cassady et al., “Network segregation varies with neural distinctiveness in sensorimotor cortex,” Neuroimage, vol. 212, no. October 2019, p. 116663, 2020, doi: 10.1016/j.neuroimage.2020.116663.

[61] Z. Zhao et al., “Altered intra- and inter-network functional coupling of resting-state networks associated with motor dysfunction in stroke,” Hum. Brain Mapp., vol. 39, no. 8, pp. 3388–3397, 2018, doi: 10.1002/hbm.24183.

[62] T. Ishida et al., “Interhemispheric disconnectivity in the sensorimotor network in bipolar disorder revealed by functional connectivity and diffusion tensor imaging analysis,” Heliyon, vol. 3, no. 6, p. e00335, 2017, doi: 10.1016/j.heliyon.2017.e00335.

[63] O. Dipasquale, L. Griffanti, M. Clerici, R. Nemni, G. Baselli, and F. Baglio, “High-dimensional ica analysis detects within-network functional connectivity damage of default-mode and sensory-motor networks in alzheimer’s disease,” Front. Hum. Neurosci., vol. 9, no. FEB, pp. 1–7, 2015, doi: 10.3389/fnhum.2015.00043.

[64] C. N. Objero, M. M. Wdowski, and M. W. Hill, “Can arm movements improve postural stability during challenging standing balance tasks?,” Gait Posture, vol. 74, no. May, pp. 71–75, 2019, doi: 10.1016/j.gaitpost.2019.08.010.

[65] O. J. Surgent, O. I. Dadalko, K. A. Pickett, and B. G. Travers, “Balance and the brain: A review of structural brain correlates of postural balance and balance training in humans,” Gait Posture, vol. 71, no. May, pp. 245–252, 2019, doi: 10.1016/j.gaitpost.2019.05.011.

[66] J. H. Jensen, J. A. Helpern, A. Ramani, H. Lu, and K. Kaczynski, “Diffusional kurtosis imaging: The quantification of non-Gaussian water diffusion by means of magnetic resonance imaging,” Magn. Reson. Med., vol. 53, no. 6, pp. 1432–1440, 2005, doi: 10.1002/mrm.20508.

[67] D. S. Novikov, J. Veraart, I. O. Jelescu, and E. Fieremans, “Rotationally-invariant mapping of scalar and orientational metrics of neuronal microstructure with diffusion MRI,” Neuroimage, vol. 174, no. March, pp. 518–538, 2018, doi: 10.1016/j.neuroimage.2018.03.006.

[68] Y. Diao and I. Jelescu, “Parameter estimation for WMTILJWatson model of white matter using encoder–decoder recurrent neural network,” Magn. Reson. Med., 2022, doi: 10.1002/mrm.29495.

[69] D. S. Novikov, E. Fieremans, S. N. Jespersen, and V. G. Kiselev, “Quantifying brain microstructure with diffusion MRI: Theory and parameter estimation,” NMR Biomed., vol. 32, no. 4, pp. 1–53, 2019, doi: 10.1002/nbm.3998.

[70] J. Cirillo, J. G. Semmler, R. A. Mooney, and W. D. Byblow, “Primary motor cortex function and motor skill acquisition: insights from threshold-hunting TMS,” Exp. Brain Res., vol. 238, no. 7–8, pp. 1745–1757, 2020, doi: 10.1007/s00221-020-05791-1.

[71] S. Papegaaij, W. Taube, M. Hogenhout, S. Baudry, and T. Hortobágyi, “Age-related decrease in motor cortical inhibition during standing under different sensory conditions,” Front. Aging Neurosci., vol. 6, no. JUN, pp. 1–8, 2014, doi: 10.3389/fnagi.2014.00126.

[72] S. Papegaaij, S. Baudry, J. Négyesi, W. Taube, and T. Hortobágyi, “Intracortical inhibition in the soleus muscle is reduced during the control of upright standing in both young and old adults,” Eur. J. Appl. Physiol., vol. 116, no. 5, pp. 959–967, 2016, doi: 10.1007/s00421-016-3354-6.

[73] I. Paparella, G. Vanderwalle, C. J. Stagg, and P. Maquet, “An integrated measure of GABA to characterize post-stroke plasticity,” NeuroImage Clin., vol. 39, no. June, p. 103463, 2023, doi: 10.1016/j.nicl.2023.103463.

[74] B. Lauber, A. Gollhofer, and W. Taube, “Differences in motor cortical control of the soleus and tibialis anterior,” J. Exp. Biol., vol. 221, no. 20, 2018, doi: 10.1242/jeb.174680.

[75] B. Lauber, A. Gollhofer, and W. Taube, “What to train first: Balance or explosive strength? Impact on performance and intracortical inhibition,” Scand. J. Med. Sci. Sport., vol. 31, no. 6, pp. 1301–1312, 2021, doi: 10.1111/sms.13939.

[76] B. Lauber and W. Taube, “Probing the link between cortical inhibitory and excitatory processes and muscle fascicle dynamics,” Sci. Rep., vol. 13, no. 1, pp. 1–13, 2023, doi: 10.1038/s41598-023-31825-z.

[77] J. T. Day, G. A. Lichtwark, and A. G. Cresswell, “Tibialis anterior muscle fascicle dynamics adequately represent postural sway during standing balance,” J. Appl. Physiol., vol. 115, no. 12, pp. 1742–1750, 2013, doi: 10.1152/japplphysiol.00517.2013.

[78] Y. A. Kuhn, S. Egger, M. Bugnon, N. Lehmann, M. Taubert, and W. Taube, “Age-related decline in GABAergic intracortical inhibition can be counteracted by long-term learning of balance skills,” J. Physiol., vol. 0, pp. 1–17, 2024, doi: 10.1113/JP285706.

[79] G. Oeltzschner et al., “Osprey: Open-source processing, reconstruction & estimation of magnetic resonance spectroscopy data,” J. Neurosci. Methods, vol. 343, no. February, p. 108827, 2020, doi: 10.1016/j.jneumeth.2020.108827.

