## Supplementary Materials for "Rebalance the inhibitory system in the elderly brain: Influence of balance learning on GABAergic inhibition and functional connectivity"

**Supplementary Methods**

**Functional MRI data processing**

First, the initial 5 time points of functional data were removed to ensure magnetic field stabilization. Functional and structural brain images were skull-stripped using SynthStrip (Hoopes et al., 2022). The skull-stripped functional images were then corrected for head motion using MCFLIRT (Jenkinson et al., 2002) implemented in FSL (<https://fsl.fmrib.ox.ac.uk/fsl/fslwiki/FSL>). Next, susceptibility distortion correction was performed using SynBOLD-Disco (Yu et al., 2023). In this pipeline, a "synthetic, undistorted" BOLD image that matches the geometry of structural T1w images and the T2* contrast was generated using a pre-trained neural network. This image was then used as reference for FSL TOPUP (Andersson et al., 2003) for distortion correction. The distortion-corrected images were then transformed to Montreal Neurological Institute (MNI)-152 2-mm space by first registering functional image to structural image with epi_reg functionality in FSL, followed by registering from structural space to MNI space using antsRegistrationSynQuick functionality implemented in Advanced Normalized Tools (ANTS) (<https://github.com/ANTsX/ANTs>). The transformed images were smoothed using a Gaussian kernel of full width at half-maximum (FWHM) of 4 mm. Next, the images in standard space were further processed with ICA-AROMA (Pruim et al., 2015), an independent component analysis (ICA) based method for removing motion-related artifacts. After this step, white matter and cerebrospinal fluid signals were regressed from the images. Finally, a high-pass filter of 0.01 Hz was applied to remove low-frequency drifts.

**Structural MRI data processing and tractography**

Following pre-processing, probabilistic streamline tractography was performed using MRtrix3 (Tournier et al., 2019). First, from the preprocessed DWI volumes, single-fiber response functions for white matter (WM), gray matter (GM), and cerebrospinal fluid (CSF) were estimated via dwi2response with the “dhollander” algorithm (Dhollander et al., 2016). Multi-shell, multi-tissue constrained spherical deconvolution was then performed to generate fiber orientation distributions (FODs) using dwi2fod (Jeurissen et al., 2014). The FOD maps were corrected for bias fields and intensity inhomogeneities via mtnormalise (Raffelt et al., 2017). Probabilistic tractography was performed to construct a “tractogram” via tckgen with the “iFOD2” algorithm (10 million streamlines, maximum tract length = 250 millimeters, FA cutoff = 0.06). The generated initial tractogram was then filtered using Spherical-deconvolution Informed Filtering of Tractograms (SIFT2) methodology. SIFT2 ensures that the (weighted) streamline reconstruction is an adequate fit of the diffusion signal at each voxel, which is itself proportional to the volume of tissue aligned in each orientation, thus providing more biologically meaningful estimates of structural connection density (Smith et al., 2015a). The left and right part of the sensorimotor network defined using group-ICA in the fMRI analysis were used to seed the streamlines, resulting mainly in streamlines passing through the anterior part of corpus callosum and connecting bilateral motor areas. Sensorimotor network structural connectivity (SNSC) was then calculated by multiplying the sum of streamline weights between bilateral nodes by the subject-specific proportionality coefficient (μ) estimated within SIFT2. This coefficient converts the values to physical units (mm^2^), making the final measure of connectivity not only proportional to cross-sectional area within a subject, but an absolute measure that is comparable across subjects (Smith et al., 2015b).

**Short interval intracortical inhibition (SICI) measurement**

Intracortical inhibition was assessed using a paired-pulse transcranial magnetic stimulation (TMS) protocol. Suprathreshold single-pulses alternated with suprathreshold pulses preceded by a subthreshold pulse (paired-pulses). The initial pulse of the paired-pulse stimulation served as a conditioning pulse, which activates intracortical inhibitory interneurons. This activation lowers the motor evoked potential (MEP) in the target muscle compared to the unconditioned stimulation (Kujirai et al., 1993; Rothwell et al., 2009). SICI was shown to be a cortical phenomenon (Di Lazzaro et al., 1998; Di Lazzaro & Rothwell, 2014; Weise et al., 2013) relying on GABA_A_ergic mechanisms (Ziemann et al., 1996). SICI measurements were performed while participants balanced on the highest achieved level of the wobble boards during performance measurements. A figure-of-eight coil (D-B80, Magventure A/S, Farum, Denmark) connected to a MagPro X100 stimulator equipped with the MagOption system (MagVenture A/S, Farum, Denmark) was used. The current was applied in an anterior-posterior direction by orienting the coil handle towards the rear. The optimal location in the left motor cortex to stimulate the right tibialis anterior (TA) muscle was defined in a seated position and marked with color on the scalp. While balancing, the coil was kept in place with a custom-made helmet, which was attached to the ceiling (Mouthon & Taube, 2019). The active motor threshold (aMT) was determined while standing and was set at the lowest intensity where three out of five consecutive stimulations produced a MEP with a peak-to-peak amplitude equal to or higher than 100 μV (Rossini et al., 2015). The initial protocol was always a test pulse intensity of 120% of the aMT with a conditioning pulse intensity of 70% of the aMT. If this protocol resulted in insufficient inhibition, the intensities of the pulses were adjusted individually. For the post-measurement the same relative intensities of the aMT were employed as for the pre-measurement, however, the aMT was re-determined in the same way as in the pre-test. The interstimulus interval for the paired-pulse stimulations was set at 2 ms (Roshan et al., 2003; Vucic et al., 2009). The stimuli were administered in two sets of 20 each, in which single- and paired-pulse stimulations were alternatively applied with an interstimulus pause of 4 seconds.

**Balance learning**

A detailed balance learning plan is provided as following.

| Exercises: week 1 | Organization | Time |
| --- | --- | --- |
| **Lesson 1: Monday - Tuesday** |  |  |
| Presentation round: first name, profession, hobby, qualifying adjective | In plenum | 10' |
| Warm up: ankle – knee – hip…  Short game: 1 – 2 – 3 (2 = touch the knee, 3 = raise the thumb) | In plenum | 10' |
| - Move the weight forward then backward: pay attention to the core - Rise on your toes, then lower on your heels | In plenum | 5' |
| Introduction of positions: one pushes the other a little   1) Natural position, each as he feels 2) Legs apart but at the same height 3) Feet tight 4) Tandem feet 5) What is the best position: ¾ 6) Turn on themselves: take the 3/4 position | By two | 10' |
| Sheathing introduction:  Lying on the back, sheathed: the other raises the legs  Standing up, sheathed, let yourself fall back: the other catches up | By two | 5' |
| **Lesson 2: Wednesday** |  |  |
| Throw a ball to one of the people by saying their first name, the person with the ball introduces themselves: first name, profession, hobby, qualifying adjective | In plenum | 10' |
| Warm up: ankle – knee – hip…  Mirror | In plenum | 10' |
| 1) Natural position, each as he feels 2) Legs apart but at the same height 3) Feet tight 4) Tandem feet 5) What is the best position: ¾   During these positions: turn the arms then turn the upper body (first without the head, then with the head) | In plenum | 10' |
| Introduction to the one-footed position   1) On one foot without moving 2) On one foot and the other forward and backward 3) Plane 4) Little ball | In plenum | 5' |
| Little fight in all positions seen |  | 10' |
| **Lesson 3: Thursday - Friday** |  |  |
| Tandem position: Throw the ball to one of the people by saying their first name, the person with the ball introduces themselves: first name, profession, hobby, qualifying adjective | In plenum | 10' |
| Warm-up: ankle – knee – hip  Resume positions on one foot  ½ turn | In plenum | 10' |
| - In pairs: eyes closed: one press turn to the left or right, two presses stop and stand on the foot on the side of the pressures - Turn the person around (eyes closed), stop then the person tries to walk straight ahead (eyes open) - ½ turn | By two | 10' |
| 1) Natural position, each as he feels 2) Legs apart but at the same height 3) Feet tight 4) Tandem feet 5) What is the best position: ¾ 6) On one foot   All positions, eyes closed. | In plenum | 10' |
| Little fight on one foot  If there is still time, a little fight on the different ones with your eyes closed (ATTENTION) only one that moves | In plenum | 5' |
| Exercises: week 2 | Organization | Time |
| **Lesson 4: Monday - Tuesday** |  |  |
| Throw a ball to one of the people by saying their first name, the person with the ball introduces themselves: first name, profession, hobby, qualifying adjective - possibly on one foot or in tandem | In plenum | 10' |
| Standard warm-up +:   - Go down to the knees then come back up and jump - Giant Steps (lunge) - One throws a tennis ball and the other tries to catch it with a cone (open side): movement - Review position on one foot | In plenum | 10' |
| - One advances in the room with his eyes closed and the other guides him only with taps on the shoulder - Turn the person around then stand on one foot | By two | 10' |
| - Face to face: try to hold the ball between the knees of the partners, one the right leg, the other the left then hold the ball between the feet, then between the feet of both partners | By two | 10' |
| 1) Two people hold a rope and the third person climbs over it on one foot 2) Small jump over the rope and land on one foot 3) On one foot make a ¼ turn then ½ turn 4) Hopscotch game | In plenum or group | 15' |
| - Little fight on one foot with theraband | By two | 5' |
| **Lesson 5: Wednesday** |  |  |
| Throw a ball to one of the people by saying their first name, the person with the ball introduces themselves: first name, profession, hobby, qualifying adjective - possibly on one foot or in tandem | In plenum | 10' |
| Standard warm-up +:   - Pantin (shift leg and arm) - On one foot and rotate the other around the body - Make the plane then the little ball | In plenum | 10' |
| Warm-up sequence: leaf – stone – scissors (or chifoumi)   - The one who loses runs and the other does squats during the time (on one foot) | By two | 10' |
| 1. One stands on two feet, eyes open at first, the partner taps on the shoulder and the person must make a quarter turn to the side of the tap. Once you understand the exercise, same thing but with your eyes closed. 2. One stands on two feet, eyes open at first and holds the rope, the other tries to unbalance him   ATTENTION: not a fight only a person who moves   - Eyes closed, two feet - Eyes closed, feet together - Eyes closed, tandem (if possible otherwise only hold in tandem with eyes closed) | By two | 10' |
| 3. Face to face: try to hold the ball between the knees of the partners, one the right leg, the other the left then hold the ball between the feet, then between the feet of both partners | By two | 5' |
| Reflex: one holds the ball up then lets it go, the other with his hands behind his back must try to catch it  Variations: change the positions of the feet (one foot, tandem, etc.) | By two | 5' |
| **Lesson 6: Thursday - Friday** |  |  |
| Throw a ball to one of the people by saying their first name, the person with the ball introduces themselves: first name, profession, hobby, qualifying adjective - possibly on one foot or in tandem | In plenum | 10' |
| Standard warm-up +:   - Make the puppet (shift legs and arms) - Mirror with me - Leg grab: trying to touch the other's thigh - Theraband: take both ends of a theraband in your hands and place one foot on the loop then push down - Theraband: hold one side in one hand and the other under the foot then with the remaining foot push to the side | In plenum | 15' |
| 1. Make a hedge with the therabands and everyone passes over them, each time standing on one foot for 2 seconds 2. Little jump over the rope as quickly as possible and when I clap my hands, get on one foot 3. Hopscotch 4. Pass the theraband (folded in 3 or 4) under the foot | In plenum | 15' |
| Reflex plus U-turn:   1) one throws the ball, the other closes his eyes: the one who throws the ball says great then throws the other opens his eyes and must catch the ball 2) same thing except that instead of closing his eyes, he turns his back and has to stand on top | By two | 10' |
| Exercises: week 3 | Organization | Time |
| **Lesson 7: Monday - Tuesday** |  |  |
| Zip game – Zap  **Rules**: the players are arranged in a circle, around the leader (a facilitator at the start of the game). The leader points to a player by saying “Zip!” ". The designated player must, as quickly as possible, give the first name of his neighbor on the right. Then the neighbor on the right must clap their hands and stand on one foot, which creates a cascade around the circle until the player designated at the base is also on one foot.  If the leader says “Zap!” », he must give the first name of his neighbor on the left and the waterfall goes to the left.  When the leader says “Zip Zap!” ", all the players change places and form a circle again, making sure that they no longer have the same neighbors.  If a player makes a mistake or does not stand on one foot, he takes the place of the leader until another player makes a mistake. | In plenum | 10 – 15’ |
| Warm-up: standard +:  1 2 3 Sun | In plenum | 10' |
| 1. In pairs: stand on one foot then throw the hoop and catch it with your hands 2. In pairs: one throws the small hoop and the other must catch it with their foot 3. Eyes open: in line on one foot: pass the hoop with your foot 4. Hold the small hoop and pass your leg through it then take your foot out 5. Place the hoop on the foot then bring the foot back while keeping the hoop hanging  - Increased difficulty: on a balance cushion | In pairs and in groups | 20' |
| A column: all on one foot: then pass a ball to each other from the sides from the front, the last one, once the ball is in hand, runs ahead and starts again. If there are enough people, two teams then in the form of a race and otherwise a column and in the form of training | In plenum | 5' |
| Final game: orbit (stand in a circle and hold hands then pass a large hoop around the circle. |  | 5' |
| **Lesson 8: Wednesday - Thursday** |  |  |
| Standard warm-up +:   - With a ball: make an 8 between the legs - Standing: raise your leg and pass the ball underneath - On one foot: eyes open and closed head tilted back - Move quickly on a line and when I clap my hands, stand on one foot - In pairs: not chased and throwing a ball at the same time | In plenum | 15' |
| Exercise repetition: throw the hoop then catch with your foot  Getting started: in the form of running: in line: passing the hoop with your foot | Two teams | 10' |
| 1) Stand on one foot, pick up a ball (small - large - heavy, etc.) according to each person's possibilities and bring it above the head, still on one foot 2) Standing on one foot, bring the free leg towards you, place the ball on the foot and keep it without dropping it 3) Standing on one foot, place the free leg at 90°, then make circles with the ball around the thigh (ball in the hands). 4) 5 people each hold a small hoop in their hands and form a line: the others go through the hoops, putting one foot in each time.   Or else   1) Back to back, on one leg, pass a ball over your head 2) Back to back, on one leg (or two if too difficult), pass the ball over your head, then between your legs 3) Back to back, on one leg (or in tandem?), pass a ball to the sides 4) 5 people each hold a small hoop in their hands and form a line: the others go through the hoops, putting one foot in each time. |  |  |
| One of the team stands at the bottom ready to catch the hoop with their foot. The next person runs, throws the hoop which the first must catch with their foot then takes their place. The first one gets in line, etc. The team that catches the most hoops wins. | Two teams |  |
| Exercises: week 4 | Organization | Time |
| **Lesson 9: Tuesday - Wednesday** |  |  |
| Standard warm-up +  In pairs: on one foot: foot against foot: try to unbalance the other (gently)  Little fight on one foot with the therabands | In plenum  By two | 10' |
| Stand facing your partner, placing a long necklace (hooked on the rings) halfway between the two of you. Bend your legs slightly, hands on your thighs. From this starting position, a third partner will give you key words (for example: head...). You will then have to touch your head with both hands. If the keyword is "knees", then you will need to touch your knees with both hands. As long as your partner tells you which body parts to touch, you will have to touch them. On the other hand, from the moment he says the word "long necklace", you will have to be the quickest to grab the long necklace hooked to the rings. => to do on one foot☺ | By three | 10' |
| **Unstable surface: 16 stack and large stack**   1) Stand on one foot then close your eyes  - Leg catcher - Then repeat the exercise of standing on one foot then closing your eyes  2) In the form of a race: passing the small hoop with your foot on the large mat 3) Work the tandem position on a large mat or 16 mat: rotate the arms + rotate the upper body 4) Little fight on one foot with theraband   Additional difficulties: close your eyes | In plenum | 20' |
| **Balance in motion: bench introduction**   - Walk on the bench right side up (just for introduction)   ***Upside down benches***   - To walk - Walk, stop in the middle and stand on one foot, then continue - Walk, stop in the middle and make a complete turn | In plenum: take the time | 10-15' |
| Final: sheathing circle: in a circle numbered 1 and 2: the body well stretched and sheathed, the 1s tilt forward and the 2 backwards all at the same time. | In plenum | 5-10' |
| **Lesson 10: Thursday - Friday** |  |  |
| Standard warm-up +  1 – 2 – 3 small goldfish (or 1 – 2 – 3 sun)   - 2nd round stop on one foot - 3rd round stop on one foot, eyes closed | In plenum | 10' |
| **Unstable surface: 16 stack and large stack**   1) Get on one foot: upper body rotation 2) On one foot: raise the leg, ¼ rotation, straighten the leg and bring it behind 3) Plane => small ball   Additional difficulties: close your eyes | In plenum | 10' |
| **Upside down benches**   1) Stand on one foot along the length of the bench 2) Get on one foot: upper body rotation 3) On one foot: raise the leg, ¼ rotation, straighten the leg and bring it behind 4) Plane => small ball 5) Stand on the bench (perpendicular to the surface), if possible on one foot 6) In pairs: back to back on a bench: pass the ball from above | In plenum | 20' |
| Two benches upside down: face to face (1.5 m)  Position yourself on the bench (on two feet) perpendicular to the surface, one part on one bench, the other on the other: quick pass with a ball   - Additional difficulty: add a smaller ball | In plenum | 10' |
| End game: day/night | In plenum | 5' |
| Exercises: 5 | Organization | Time |
| **Lesson 11: Monday - Tuesday** |  |  |
| Standard warm-up +  Contacting Swissball   - Tilt the pelvis forwards and backwards then to the sides, then rotation - Move forward while sitting on the swissball (small jump) - Sit: raise one foot then the other - In pairs back to back with a swissball in between: go down and up (squats) | In plenum | 15' |
| Game: Modified burnt ball: the running team starts, those in the middle run to the bench upside down (one of the two sides) and all stand on it, when everyone is on it (without falling), the person running must be on a base otherwise they get burned. | In plenum | 10' |
| **Unstable surfaces**  Large mat – mat of 16   - Run in place then at the stop sign get on one foot - Stand on one foot and tilt your head back - Walk with your eyes closed on the big carpets (in pairs) | In plenum | 10' |
| **Upside down benches**:   - Walk on the bench over obstacles (some participants hold a stick and adjust the height) - Step on the bench and in the middle, squat - Additional difficulties: teapot - No giants - In pairs: face to face: try to hold a small ball between the foot then between the knees - In pairs: small fights on two feet - Additional difficulties: on one foot (if too great a difference in level between two, one on one foot and the other on both. | In plenum | 20' |
| Final challenge: try to cross each other on the bench upside down | By two | 5' |
| **Lesson 12: Wednesday** |  |  |
| Standard warm-up +   - Sitting on the swissball: one throws the small hoop the other must catch it with one foot sitting on the swissball (the one throwing is standing) - Swissball: try to sit without your feet on the ground | In plenum | 15' |
| **From the bench upside down**:   - Jump on a 16 mat with balanced landing (stable) - Same with ¼ - ½ or 1 full turn - Same thing with landing on one foot - Same thing with eyes closed (on two feet and if too easy on one foot) | In plenum | 15' |
| **Upside down benches**:   - From a top of a box: jump onto the bench upside down - Two benches facing each other: each on a bench:   badminton   - Moving in a shuffled step on the bench | In plenum | 15' |
| Mikado: two upside down benches: at the end a box with two mikados   - Go to the bench and take a mikado, get off the bench, run around a pole and return to the starting line. When the first has got off the box, the second can leave. The team with the most sticks wins. | In plenum | 10' |
| **Lesson 13: Thursday - Friday** |  |  |
| Standard warm-up  + sit on a swissball without your feet | By two | 15' |
| Stand facing your partner, placing a long necklace (hooked on the rings) halfway between the two of you. Bend your legs slightly, hands on your thighs. From this starting position, a third partner will give you key words (for example: head...). You will then have to touch your head with both hands. If the keyword is "knees", then you will need to touch your knees with both hands. As long as your partner tells you which body parts to touch, you will have to touch them. On the other hand, from the moment he says the word "long necklace", you will have to be the quickest to grab the long necklace hooked to the rings. => to do on a bench then on one foot on the bench | By three | 10' |
| Upside down bench   - Bench – box – bench – box – bench: be careful to secure the bench that goes up and down either with a top of the box or against the wall - Normal passage - Stop in the middle: make a half or full turn - Get on one foot | In plenum | 15' |
| Upside down bench   - One on the bench: the other on the ground throws him a ball but causes movements - Standing on the bench   => Theraband: take both ends of a theraband in your hands and place one foot on the loop then push down  => Theraband: hold one side in one hand and the other under the foot then with the remaining foot push to the side  => Hold both ends of the theraband in your hands and pass your leg over them | By two | 15' |
| Finale: little fight on the benches with therabands | By two | 10' |
| Exercises: 6 | Organization | Time |
| **Lesson 15: Monday - Tuesday** |  |  |
| Standard warm-up +   - Pedalo - Small in-line rocking board (first offset then aligned) - Swissball | In plenum / in small groups | 15' |
| 1 – 2 – 3 sun sitting on the swissball | In plenum | 5-10' |
| Unstable surface: 16 mat or large mat   - One on the unstable surface must catch with their foot the small hoop thrown by the other | By two | 5' |
| Upside down bench   - Start by simply walking on it, stop in the middle, stand on one foot, go down... - Walk while passing yourself with a small ball - Walk with your leg raised and pass the ball - Same thing as on the unstable surface: one throws the small hoop, the other must catch it with their foot - Online passage of the small hoop from one foot to the other (if works well in running form)   => repeat before once on the ground | In plenum | 15' |
| Two benches one behind the other: participants pass over the two benches upside down at the end of the benches, they must take a hoop and throw it at one of the three stakes (1 stake = 1 point / 2nd = 2 points / 3rd = 3 points) | By team | 10' |
| Swissball: sitting and lying on your stomach |  | 10' |
| **Lesson 16: Wednesday** |  |  |
| Standard warm-up +   - Pedalo - Mexican hat – roll – rolabola - Swiss ball | In plenum | 15' |
| Estafette: two teams: 3 poles – half of the group on one side, the other on the other side   - Run to the middle pole, turn around 4 x then continue to the next pole to hand over to the next one | By teams | 10' |
| Upside down bench on the ground in front of the rings, bar fixed on the rings: sitting on the bar, start from a box and land on an upside down bench:   - Direction of length in relation to the rings: advance to the end of the bench - Direction of width (vertical) in relation to the rings   Try without holding on to the rings | In two groups  Alternation with the next block | 10' |
| Benches backwards:   - Passage on the benches, eyes closed - Several benches upside down position parallel: go over the series of benches, stopping on each bench (spread the benches apart to increase the difficulty) - Rocker (bench on top of box) | In two groups  Alternation with the front block | 10' |
| Swissball on your knees |  | 10' |
| Handkerchief: in a circle all on one foot (with one then with two handkerchiefs) | In plenum | 5' |
| **Lesson 17: Thursday - Friday** |  |  |
| Standard warm-up +   - Pedalo - Mexican hat – roll – rolabola - swissball - Introduction to small carpets (getting pulled) |  | 20' |
| Tail catcher: jumper hung in training |  | 10' |
| 1) Benches attached to the wall bars: stand on two small carpets and descend (firstly, descend seated) 2) Flying carpet (top of box with basketball underneath) | Two groups  Alternating 1 and 2 | 10' |
| Reverse benches: reflex exercise   - In pairs: one lets go of the stick (with one or two), the other must catch it with their hands behind their back. - In pairs: one closes their eyes, the other says great then throws a small (soft) ball |  | 10' |
| Upside down bench   - Half turn with feet apart - Not chased - Face to face on the bench, palm to palm, a complete turn and we join palm to palm |  | 10' |
| Two parallel benches, each team on a bench in a column: passing the ball from above, then from the side. The ball starts in front and finishes behind, the last one gets off the bench and runs to stand in front.  First team to finish wins | By team | 5' |
| Exercises: 7 | Organization | Time |
| **Lesson 18: Monday - Tuesday** |  |  |
| Standard warm-up +  Swissball strengthening   - On one foot, roll the swissball around you - In pairs back to back, wedge the swissball between the two backs then lower into the knees => for those who manage to sit on the ground and get up | In plenum | 20' |
| Handkerchief: those in the circle stand on one foot | In plenum | 10' |
| Swissball:   - Sat - Sit on the swissball then put your feet on a volleyball or basketball - In threes: sit on the swissball then place your feet on the swissball of the person in front (whose feet are on the ground). The third person for belaying. - Facing each other: passing with a ball (without putting your feet on the ground) - Small fights with theraband: one with his feet on the ground and not the other - Then the two without their feet on the ground | By two or three | 25' |
| Swissball: bench   - Walk on the bench upside down, holding the swissball at arm's length above your head - Stand on one foot and place the swissball to the right then to the left - Same as the warm-up: back to back, swissball stuck between the two backs and down into the knees | In plenum | 10' |
| - Lying down: let go of one hand then the other then both - Kneeling |  | 10' |
| Final challenge:WARNING: difficult, only those who want, others can continue seated | In plenum | 5' |
| **Lesson 19: Wednesday** |  |  |
| Standard warm-up +  Balance mat on one leg | In plenum | 10' |
| Game: tic tac toe (hoop + cone) | By team | 10' |
| Bench backwards:   - Hold a small ball between the knees then between the feet - Standing on the bench   => Theraband: take both ends of a theraband in your hands and place one foot on the loop then push down  => Theraband: hold one side in one hand and the other under the foot then with the remaining foot push to the side  => Hold both ends of the theraband in your hands and pass your leg over them   - Walk on the bench upside down, holding the swissball at arm's length above your head - Stand on one foot and place the swissball to the right then to the left - Same as the warm-up: back to back, swissball stuck between the two backs and down into the knees | By two  In plenum | 20' |
| - First sit on the bar in the middle of the rings in balance, if stable swing slightly - Same thing but swing harder (put big stacks under) | In plenum | 10' |
| - Badminton on the benches |  | 10' |
| Orbit on the benches in racing form | By team | 5' |
| **Lesson 20: Thursday - Friday** |  |  |
| Warm-up +   - Mexican hat - Rolabola - Pedalo - Swissball | In plenum | 10' |
| Upside down bench   - Spin sumo on the bench - Not chased - In pairs: one throws the ball and the other catches it - First with the hands then with the foot - In pairs: one throws a hoop and the other tries to catch it with their foot - Jump from the top of the box onto the bench | In plenum | 10' |
| Estafette memory: run over a large mat, then on the ground before getting on an upside-down bench and turning over two memory cards (if family I take otherwise I leave) | As a team | 10' |
| Swing from the rings and land on a balance cushion or a 16 mat or a large mat: on two feet then on one foot | In plenum | 10' |
| Challenge: make a circle all seated on the swissball and put their feet on the ball of the one in front (no one must have their feet on the ground) | In plenum | 10' |
| Exercises: 8 | Organization | Time |
| **Lesson 21: Monday - Tuesday** |  |  |
| Standard warm-up  + in pairs: passing a ball to each other while moving in a shuffle (one leads, the other follows)  + Swissball: sit on the ball and move forward making small hops  + Swissball: sit on it | In plenum | 20' |
| Game: Horse racing: two teams: one circle inside and the other outside: the inside team stands on one foot passes a volleyball meanwhile, the team on the The outside runs around the circle 3 times! The team that has completed the most turns with the balloons is the winner! (ATTENTION: part of the warm-up, so easy running not sprinting) | By team | 10' |
| Training :  Take the different elements of Wednesday's course over the two days and train them  **Monday: 1-6-7**  **Tuesday: 3 – 5 – 8** | Per station: 7' per station per group | 20' |
| Challenge: make a circle all seated on the swissball and put their feet on the ball of the one in front (no one must have their feet on the ground)  Monday: just in twos or threes  Tuesday: a full circle (two teams): one team does, the other ensures | By team (plenum) | 10' |
| **Lesson 22: Wednesday – Thursday evening** |  |  |
| Standard warm-up  + tandem position: turn the upper body  + position on one foot: turn the upper body |  | 10' |
| Scrabble game: the team stays around a “table” and one after the other runs to look for a letter (one letter at a time). Meanwhile the rest of the team tries to make as many words as possible. |  | 10' |
| Course :   1) Start: rings with bar in the middle landing on the bench upside down, lengthwise - Walk along the entire bench 2) Slalom between the poles 3) Bench – box – bench attached to the wall bars – box – two benches attached to the wall bars large mats underneath (walk on it – go down like skiing) 4) Swissball: sit on the ball and move forward making small hops 5) Reverse bench: move forward and the other passes with the ball 6) Pedalo 7) Go over 3 boxes (not at the top and quite close) then jump onto a large carpet 8) Passing hurdles: while walking and always take a break on one foot between two | The entire route is done in pairs | 30' |
| Game: reflex circle |  | 5' |
| **Lesson 23: Thursday morning - Friday** |  |  |
| Standard warm-up  + Reflex game:   - One stands behind him, the other throws and says great when he throws the little ball, the first must catch the little ball - One stands on one foot, eyes closed, the other throws and says great as he throws the small ball, the first must catch the ball | In plenum | 15' |
| Estafette: two teams: passage on a bench, bench fixed to the box (not too stiff), on the box pieces of paper with numbers.  Everyone must run on the course to turn a number, remember the number, at the end the whole team calculates the sum of all the numbers. Once the sum is done, the team shouts out the number. | By team  Prepare the numbers on small pieces of paper | 10' |
| Benches backwards:   - One sits on the bench and tries to return the volleyball thrown by the second with his foot. - Badminton on the benches (facing each other) both lie on the bench - One stands on two feet (too easy on one foot), eyes closed, the other throws and says top as he throws the little ball, the first must catch the ball - Passing the small hoop over the foot | In plenum | 20' |
| Race: two teams or more if many people in a column on an upside-down bench   - Passing the ball from above from the front, the last one comes down from the bench running in front, gets back on the bench and passes the ball again (all the others must take a step back to make room): the first team whose members are all went ahead and won | By team | 10' |
| Exercises: week 9 | Organization | Time |
| **Lesson 24: Monday and Tuesday** |  |  |
| Standard warm-up  + small rolled green mat: on one foot (eyes open/closed) – tandem position: turn the upper body |  | 10' |
| Upside down benches   - One on the ground throws a small ball and on the bench the other must catch it with a cone - One on the ground throws a volleyball and the other on the bench must throw it with his foot - Badminton | By three | 20' |
| Scrabble: upside down bench + table box (one after class looks for a letter, the others try to make as many points as possible with words (no first name or city name, etc.) | 3 – 4 teams | 5' |
| Stand facing your partner, placing a long necklace (hooked on the rings) halfway between the two of you. Bend your legs slightly, hands on your thighs. From this starting position, a third partner will give you key words (for example: head...). You will then have to touch your head with both hands. If the keyword is "knees", then you will need to touch your knees with both hands. As long as your partner tells you which body parts to touch, you will have to touch them. On the other hand, from the moment he says the word "long necklace", you will have to be the quickest to grab the long necklace hooked to the rings. => on the benches | By 3 | 10' |
| Bar on the rings: one side of the bench in the place fixed on the bar: up-down | By group on the ring slopes | 10' |
| **Lesson 25: Wednesday and Friday (Thursday: ascent)** |  |  |
| Standard warm-up  + swissball:   - On one foot + tandem position: place the ball on R then on L - On one foot + tandem position: turn - Sitting position   => lift one foot then the other  => put one foot on the other knee  => without feet | Take some time with the swissballs | 20' |
| Bench on unstable surface   - Bar on the rings: one side of the bench in the place fixed on the bar: up-down - Bench upside down on one side on a small mat - Bench upside down on a 16 mat - Cross the benches, make a complete turn, stand on one foot… | In plenum (3 alternating groups) = 7' per station | 25' |
| Game: Pass the ball from above from the front, the last course in front and passes the ball again and so on: in the form of a race | In plenum | 10' |
| If you still have time, take up swissball for a while but take it easy | In plenum | 5' |
| Exercises: 10 | Organization | Time |
| **Lesson 26: Monday and Wednesday** |  |  |
| Standard warm-up  + Large rug:   - Run in place then at my signal get on one foot - Thigh catcher - medicine ball: hold on to it (WARNING in pairs: especially when going up) | In plenum | 20' |
| Warm-up: 1 – 2 – 3 sun: stand on one foot without moving | In plenum | 10' |
| To begin with :   - Bar on the rings: one side of the bench in the place fix on the bar: go up-down: put them at ease - In the form of a competition (NOT THE SPEED THAT COUNTS BUT THE NUMBER OF BALLS IN THE BOX AND THE NUMBER OF PINS AT THE BOTTOM) - Two teams: short race on a short stretch, climb on the bench, aim with a ball in a box (without the top) then go back down (WARNING: the goal is not to go fast but to aim well without holding on to the ropes of the rings) the team having put the most balls in the box wins (2 – 3 balls per person depending on the number of participants) - Bench upside down on a mat of 16 (or only one side above): shot with a volleyball on 3 or 4 pins on a box and add the number of pins reversed (if no pins you can put on ball on a cone) | In plenum then by team | 20' |
| Reflex circle | In plenum (or two groups | 5' |
| **Lesson 27: Tuesday and Thursday** |  |  |
| Standard warm-up  + Mexican hat  +Rolabola  + Pedal boat  + Balance mat… | In plenum | 15' |
| Seated balance (gain)   - Sitting on a swissball, feet either on a bottom of the box, then on a medicine ball then on a volleyball/basketball then on another swissball - Bench attached to the wall bars: descents seated on a carpet without touching with your feet (optional: at the same time aim at the stakes with small hoops) - Bench suspended from the rings in the right place: sit on one and the other moves | In plenum (depending on the number of participants 3 alternating groups, 7' per station) | 25' |
| Game: tail catch | In plenum | 5' |
| Balance (repetitions)   - Bar on the rings: one side of the bench in the place fixed on the bar: up-down - Bench upside down on one side on a small mat - Bench upside down on a 16 mat - Cross the benches, make a complete turn, stand on one foot… | In plenum | 15' |
| **Lesson 28: Friday** |  |  |
| Warming up  + in a circle, pass each other with a volleyball as quickly as possible (in groups if too many participants), always staying in motion  + same thing on one foot  + passing the ball with the foot (football) | In plenum | 15' |
| Game: two teams or three   - Bar on the rings: one side of the bench in the place fixed on the bar: up-down - At the end of each bench, a box with a mikado on it | By team | 10' |
| Bar on the rings: one side of the bench at the place fixed on the bar: go up-down (put the rings low enough)   - When you reach the top, get on one foot - Arrived at the top, on two feet, eyes closed | In plenum | 15' |
| Bar on the rings: fixed bench upside down on top   - Go up go down - Arrived at the top, stand on one foot (to see if possible)   If many participants: install a bench upside down on two small mats at each end | In plenum | 15' |
| Reflex circle if time | In plenum | 5' |
| Exercises: week 11 | Organization | Time |
| **Lesson 27: Tuesday and Friday** |  |  |
| Standard warm-up  + Mexican hat  + small board that rocks |  | 15' |
| Bar on the rings: fixed bench upside down on top   - Go up go down - Arrived at the top, stand on one foot (to see if possible)   (if too easy you can put a small mat under the side which is on the ground and it moves even more) | In plenum | 20' |
| 3 positions:   - Bench upside down on one side on a small mat - Bench upside down on a 16 mat - Cross the benches, make a complete turn, stand on one foot… - Reck bar (fixed at the lowest): walk on it, stand on one foot | 7 min per shift | 20' |
| Game: day / night |  | 5' |
| **Lesson 28: Wednesday and Thursday** |  |  |
| Standard warm-up |  | 5' |
| 1. Box top on basketballs 2. Bench hang on the rings (right side up): go up to where you feel comfortable then stand on the side and a third person moves the side of the rings 3. Reverse bench on sticks: move back and forth 4. Bench upside down on casters: slalom between poles 5. Empty the carpet carts: one standing then the other moving in all directions | 7' per post | 40' |
| In a relaxed way to have fun  + one throws a small ball the other catches with a cone (on the ground then on a bench upside down)  + badminton (on the ground then on the benches) |  | 15' |
| Exercises: 12 | Organization | Time |
| **Lesson 29: Monday and Wednesday** |  |  |
| Standard warm-up  + In pairs, one walks in a straight line on the ground, the other on the side holds a swissball and lightly pushes the one walking to destabilize him.   - Also with eyes closed | In plenum | 15' |
| Bar on the rings: fixed bench upside down on top   - Go up go down - Arrived at the top, stand on one foot (to see if possible)   (if too easy you can put a small mat under the side which is on the ground and it moves even more)  **Those who are not busy: Mexican hat** | In plenum: make two tracks | 15' |
| Tail catcher: to relax |  | 5' |
| Upside down bench on sticks   - In the direction of the bench: two on top and each moves the bench (kind of a little fight) ATTENTION: put yourself between the sticks - Position yourself perpendicular to the bench and another moves the bench (being at the bottom of the bench) - One throws a volleyball and a small ball (always a little changed) and the other on the bench perpendicularly catches. | In plenum: make two or 3 tracks (depending on the number of sticks available)  If there is not enough bench, do not hesitate to do the same thing but on a bench that does not move (for those who are not sure. | 20' |
| On the bench: small fight in the form of a game so relax with swissball (each holds a swissball or both hold one) then try to unbalance (you have to try on the ground first if each takes a swissball so that they understand that 'you have to go gently) | By two + two who secure | 5' |
| **Lesson 30: Tuesday and Friday** |  |  |
| Standard warm-up  + on bench upside down  1st task: count to 3, repeat faster and faster  2nd task: raise the left hand in place of the 2  3rd task: touch your right knee with your left hand  4th task return to 1st task |  | 15' |
| Bench on casters:   - On two feet: move right – left - On one foot: quietly move forward   Carpet cart:   - On two feet: move - Then on one foot   **Those who are not busy: Mexican hat** | By alternating group | 20' |
| Game: the circle: one person in the middle: the others try to change places by making a discreet sign and the one in the middle must try to take the place in the circle |  | 10' |
| Bench upside down on a small mat (on one or both sides)   - Cross the benches, make a complete turn, stand on one foot…   Upside down bench on two small mats   - To walk on - Stop in the middle stand on one foot - Stop in the middle and stand perpendicular - One on the bench and the other on the ground passes to him (without moving) | By alternating group | 20' |
| Exercises: 13 | Organization | Time |
| **Lesson 31: Monday and Wednesday** |  |  |
| Standard warm-up  + passage on the benches with counting backwards  + Mexican hat  + board |  | 20' |
| Game: catch tail by team (blue against red for example) |  |  |
| Bench upside down on a small mat (on one or both sides)   - Cross the benches, make a complete turn, stand on one foot…   Bench upside down on two small mats (one on each side)   - To walk on - Stop in the middle stand on one foot - Giant step - Airplane – small ball - Moving in chased steps - Stop in the middle and stand perpendicular - One on the bench and the other on the ground passes him with a volleyball (without moving) | By alternating group | 25' |
| Little theraband fight on the moving benches |  | 10' |
| **Lesson 32: Tuesday and Thursday** |  |  |
| Standard warm-up  + giant steps  + throw a volleyball: important to look for lateral movement (not chased)  + one throws the ball and the other has his back turned: the first says up and throws the ball and the second turns around and catches the ball  + Mexican hat |  | 20' |
| One closes his eyes and the other turns it on itself (4-5 times) then stops it: the other must:   - Walk straight - The moment he stops it, open your eyes and get on one foot |  | 10' |
| Bench upside down on two small mats (one on each side) or one side:   - Not chased - Throw a small ball and the other catches it with an upside down cone - Obstacle crossing (holding a hurdle bar) | By alternating group | 25' |
| Jump from the top of a box onto an upside-down bench or from the mats (but the mat is soft and therefore more difficult) and stabilize on the band |  |  |
| Game: day/night |  |  |
| **Lesson 33: Friday** |  |  |
| Standard warm-up  + on one foot: pick up balls of different sizes and weights then bring them back above your head  + Mexican hat |  | 25' |
| Game: day/night with ball (PLEASE NOTE: please note that this is still a warm-up)   - Each one a small ball: if you say day, day throw the ball to night and run away and night catches the ball and tries to catch day and vice versa. |  | 10' |
| On the moving bench: either a small mat underneath on one side, or on both sides:  Estafette per team (2 teams):  Run one end, cross a bench which is on two small mats, slalom between pole, bench box bench, return passage on a large mat and give the relay to the next one. | By alternating group | 10' |
| On the moving bench  - Theraband: take both ends of a theraband in your hands and place one foot on the loop then push down  - Theraband: hold one side in one hand and the other under the foot then with the remaining foot push to the side  - Hold both ends of the theraband in your hands and pass your leg over them |  | 15' |
| Passing the small hoop on the bench (without it moving) in the form of a race |  | 5' |
| Exercises: 14 | Organization | Time |
| **Lesson 34: Monday and Wednesday** | | |
| Standard warm-up  + Mexican hat  + boards | Groups for Mexican hats and planchettes | 15' |
| One closes his eyes and the other turns it on itself (4-5 times) then stops it: the other must:   - Walk straight - The moment he stops it, open your eyes and get on one foot | By 2 | 10' |
| Upside down benches on a small mat (on one side)   - Walk normal - Make a complete turn in the middle of the bench - Get on one foot | 1 walking on the bench, 1 on each side to provide security just in case, 1 on the mat towards the end of the bench to stabilize if the bench moves too much => ideally groups of 4 | 20' |
| Estafette with benches upside down on a small carpet on one side. 1-2m behind the carpet, place a box without the top   - Cross the bench with a volleyball in your hands, at the end of the bench try to throw the ball into the box.   Inside = 1 pt  Outside = 0 pt | By team  Not a race, what counts is the number of points scored. For example, do 2 passes per person.  Belay as in the previous exercise. | 15' |
| **Lesson 35: Tuesday and Thursday** | | |
| Standard warm-up  + plane  + small ball  + reflex circle (in a circle, everyone has a stick, say “left” or “right” and everyone moves to the right side to grab the stick that is there) | In plenum | 15' |
| 3 upside down benches in a line with box top between two (continuous line that participants can do without getting off)   - Cross normal - Stop in the middle of each bench   - Stand on one foot   - Make the little ball   - Make the plane - Cross counting backwards | 1 person walking on the bench and 1 on each side to provide security just in case | 20' |
| Bench upside down on two small mats (one on each side)   - Cross normal - Obstacle crossing (holding a hurdle bar) - Moving in chased steps - Moving in a shuffled step, in the middle throw a volleyball into a box (put a box without the top not too far from the bench) | 1 walking on the bench, 1 on each side to provide security just in case, 1 towards one end of the bench to stabilize if the bench moves too much | 20' |
| Small theraband fight on benches in reverse: 2 people on the bench, they hold the theraband together (the middle with both hands, and the other the two ends), try to destabilize the other by pulling / moving the theraband | People on the sides to provide insurance just in case | 10' |

**Sample size calculation**

The required number of participants was estimated based on an a priori power analyses (Faul et al., 2007) for a repeated measures ANOVA with the following assumptions: effect size 0.25, alpha error probability 0.05, power 95 %, and correlation among repeated measures 0.8. The power analyses revealed a total sample size of 24 participants (12 per group). The effect size of 0.25 was chosen based on our just published results (Kuhn et al., 2024) on balance training and SICI, where an effect size of 0.77 was reported for SICI and an effect size of 0.5 was reported for balance performance. Thus, our choice of effect size is reasonable.

The power analyses revealed a total sample size of 24 participants (12 per group). Considering the challenge of our longitudinal study with physical training and to be sure to reach the statistical power needed, we calculated the effective sample size by a high estimated dropout rate (note that the drop out at either pre- or post-measurement will double the rate, because a complete data requires both pre- and post-measurements), leading to 40 participants in total, i.e., 20 participants in each group.”

**Number of statistical tests**

In this study we performed the following statistical tests:

| Description | Number of tests performed | Correction |
| --- | --- | --- |
| Linear mixed-effect modelling for output parameters (SICI, GABA, Performance, VSN-FC, DSN-FC, SNSC) | 6 | Hypothesis driven; no correction performed |
| Two-sample t-tests to determine direction of change | 2 for each mixed-effect analysis | Corrected using Benjamini-Hochberg procedure |
| Correlation analysis – changes and baseline | 5 in learning group, 5 in both groups | Corrected using Benjamini-Hochberg procedure |

**References**

Andersson, J. L. R., Skare, S., & Ashburner, J. (2003). How to correct susceptibility distortions in spin-echo echo-planar images: Application to diffusion tensor imaging. *NeuroImage*, *20*(2), 870–888. https://doi.org/10.1016/S1053-8119(03)00336-7

Dhollander, T., Raffelt, D., & Connelly, A. (2016). Unsupervised 3-tissue response function estimation from single-shell or multi-shell diffusion MR data without a co-registered T1 image. *ISMRM Workshop on Breaking the Barriers of Diffusion MRI*, *September*, 1–2. https://www.researchgate.net/publication/307863133

Di Lazzaro, V., Restuccia, D., Oliviero, A., Profice, P., Ferrara, L., Insola, A., Mazzone, P., Tonali, P., & Rothwell, J. C. (1998). Magnetic transcranial stimulation at intensities below active motor threshold activates intracortical inhibitory circuits. *Experimental Brain Research*, *119*(2), 265–268. https://doi.org/10.1007/s002210050341

Di Lazzaro, V., & Rothwell, J. C. (2014). Corticospinal activity evoked and modulated by non-invasive stimulation of the intact human motor cortex. *Journal of Physiology*, *592*(19), 4115–4128. https://doi.org/10.1113/jphysiol.2014.274316

Faul, F., Erdfelder, E., Lang, A.-G., & Buchner, A. (2007). G*Power 3: A flexible statistical power analysis program for the social, behavioral, and biomedical sciences. *Behavior Research Methods*, *39*(2), 175–191.

Hoopes, A., Mora, J. S., Dalca, A. V., Fischl, B., & Hoffmann, M. (2022). SynthStrip: skull-stripping for any brain image. *NeuroImage*, *260*(March), 119474. https://doi.org/10.1016/j.neuroimage.2022.119474

Jenkinson, M., Bannister, P., Brady, M., & Smith, S. (2002). Improved Optimization for the Robust and Accurate Linear Registration and Motion Correction of Brain Images. *NeuroImage*, *17*(2), 825–841. https://doi.org/10.1006/nimg.2002.1132

Jeurissen, B., Tournier, J. D., Dhollander, T., Connelly, A., & Sijbers, J. (2014). Multi-tissue constrained spherical deconvolution for improved analysis of multi-shell diffusion MRI data. *NeuroImage*, *103*, 411–426. https://doi.org/10.1016/j.neuroimage.2014.07.061

Kuhn, Y. A., Egger, S., Bugnon, M., Lehmann, N., Taubert, M., & Taube, W. (2024). Age-related decline in GABAergic intracortical inhibition can be counteracted by long-term learning of balance skills. *Journal of Physiology*, *0*, 1–17. https://doi.org/10.1113/JP285706

Kujirai, T., Caramia, M. D., Rothwell, J. C., Day, B. L., Thompson, P. D., Ferbert, A., ..., & Marsden, C. D. (1993). Corticocortical inhibition in human motor cortex. *Journal of Physiology*, *471*(1), 501–519. https://doi.org/10.1016/0006-2952(95)00035-X

Mouthon, A., & Taube, W. (2019). Intracortical Inhibition Increases during Postural Task Execution in Response to Balance Training. *Neuroscience*, *401*, 35–42. https://doi.org/10.1016/j.neuroscience.2019.01.007

Pruim, R. H. R., Mennes, M., van Rooij, D., Llera, A., Buitelaar, J. K., & Beckmann, C. F. (2015). ICA-AROMA: A robust ICA-based strategy for removing motion artifacts from fMRI data. *NeuroImage*, *112*, 267–277. https://doi.org/10.1016/j.neuroimage.2015.02.064

Raffelt, D., Tournier, J.-D., Smith, R. E., Dhollander, T., Tournier, J.-D., Tabbara, R., Smith, R. E., Pierre, E., & Connelly, A. (2017). Bias Field Correction and Intensity Normalisation for Quantitative Analysis of Apparent Fibre Density. *Proceedings of the International Society for Magnetic Resonance in Medicine*, *April*, 3541. https://www.researchgate.net/publication/315836355

Roshan, L., Paradiso, G. O., & Chen, R. (2003). Two phases of short-interval intracortical inhibition. *Experimental Brain Research*, *151*(3), 330–337. https://doi.org/10.1007/s00221-003-1502-9

Rossini, P. M., Burke, D., Chen, R., Cohen, L. G., Daskalakis, Z., Di Iorio, R., Di Lazzaro, V., Ferreri, F., Fitzgerald, P. B., George, M. S., Hallett, M., Lefaucheur, J. P., Langguth, B., Matsumoto, H., Miniussi, C., Nitsche, M. A., Pascual-Leone, A., Paulus, W., Rossi, S., … Ziemann, U. (2015). Non-invasive electrical and magnetic stimulation of the brain, spinal cord, roots and peripheral nerves: Basic principles and procedures for routine clinical and research application: An updated report from an I.F.C.N. Committee. *Clinical Neurophysiology*, *126*(6), 1071–1107. https://doi.org/10.1016/j.clinph.2015.02.001

Rothwell, J. C., Day, B. L., Thompson, P. D., & Kujirai, T. (2009). Short latency intracortical inhibition: One of the most popular tools in human motor neurophysiology. *Journal of Physiology*, *587*(1), 11–12. https://doi.org/10.1113/jphysiol.2008.162461

Smith, R. E., Tournier, J. D., Calamante, F., & Connelly, A. (2015a). SIFT2: Enabling dense quantitative assessment of brain white matter connectivity using streamlines tractography. *NeuroImage*, *119*, 338–351. https://doi.org/10.1016/j.neuroimage.2015.06.092

Smith, R. E., Tournier, J. D., Calamante, F., & Connelly, A. (2015b). The effects of SIFT on the reproducibility and biological accuracy of the structural connectome. *NeuroImage*, *104*, 253–265. https://doi.org/10.1016/j.neuroimage.2014.10.004

Tournier, J. D., Smith, R., Raffelt, D., Tabbara, R., Dhollander, T., Pietsch, M., Christiaens, D., Jeurissen, B., Yeh, C. H., & Connelly, A. (2019). MRtrix3: A fast, flexible and open software framework for medical image processing and visualisation. *NeuroImage*, *202*(January), 116137. https://doi.org/10.1016/j.neuroimage.2019.116137

Vucic, S., Cheah, B. C., Krishnan, A. V., Burke, D., & Kiernan, M. C. (2009). The effects of alterations in conditioning stimulus intensity on short interval intracortical inhibition. *Brain Research*, *1273*, 39–47. https://doi.org/10.1016/j.brainres.2009.03.043

Weise, D., Mann, J., Ridding, M., Eskandar, K., Huss, M., Rumpf, J. J., Di Lazzaro, V., Mazzone, P., Ranieri, F., & Classen, J. (2013). Microcircuit mechanisms involved in paired associative stimulation-induced depression of corticospinal excitability. *Journal of Physiology*, *591*(19), 4903–4920. https://doi.org/10.1113/jphysiol.2013.253989

Yu, T., Cai, L. Y., Morgan, V. L., Goodale, S. E., Englot, D. J., Chang, C. E., Landman, B. A., & Schilling, K. G. (2023). SynBOLD-DisCo: synthetic BOLD images for distortion correction of fMRI without additional calibration scans. *Proceedings of SPIE--the International Society for Optical Engineering*, *12464*. https://doi.org/10.1117/12.2653647

Ziemann, U., Rothwell, J. C., & Ridding, M. C. (1996). Interaction between intracortical inhibition and facilitation in human motor cortex. *Journal of Physiology*, *496*(3), 873–881. https://doi.org/10.1113/jphysiol.1996.sp021734

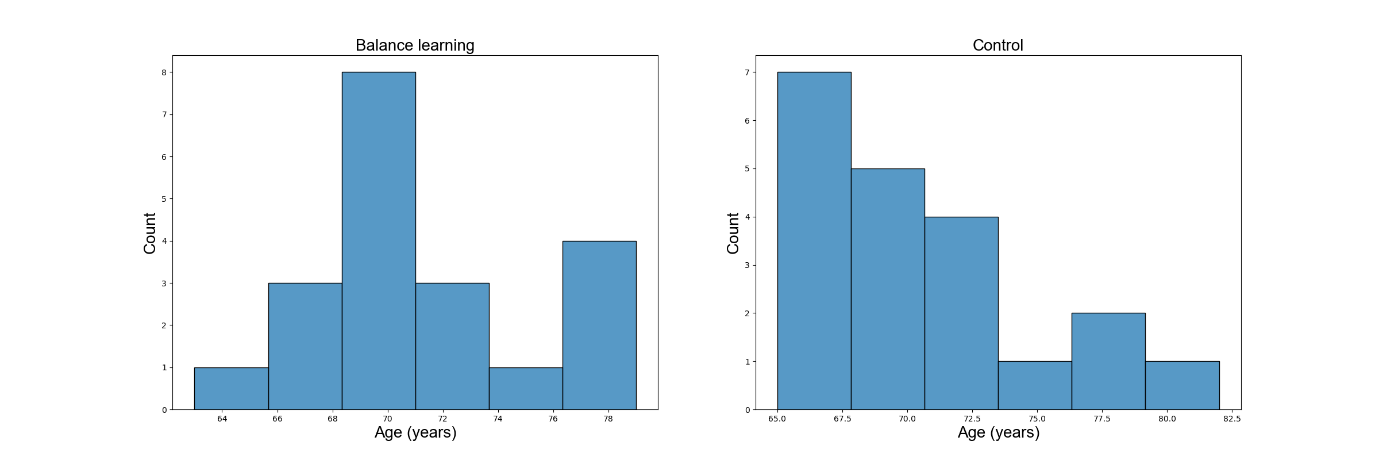

Figure 1. Age distribution of participants in both groups.

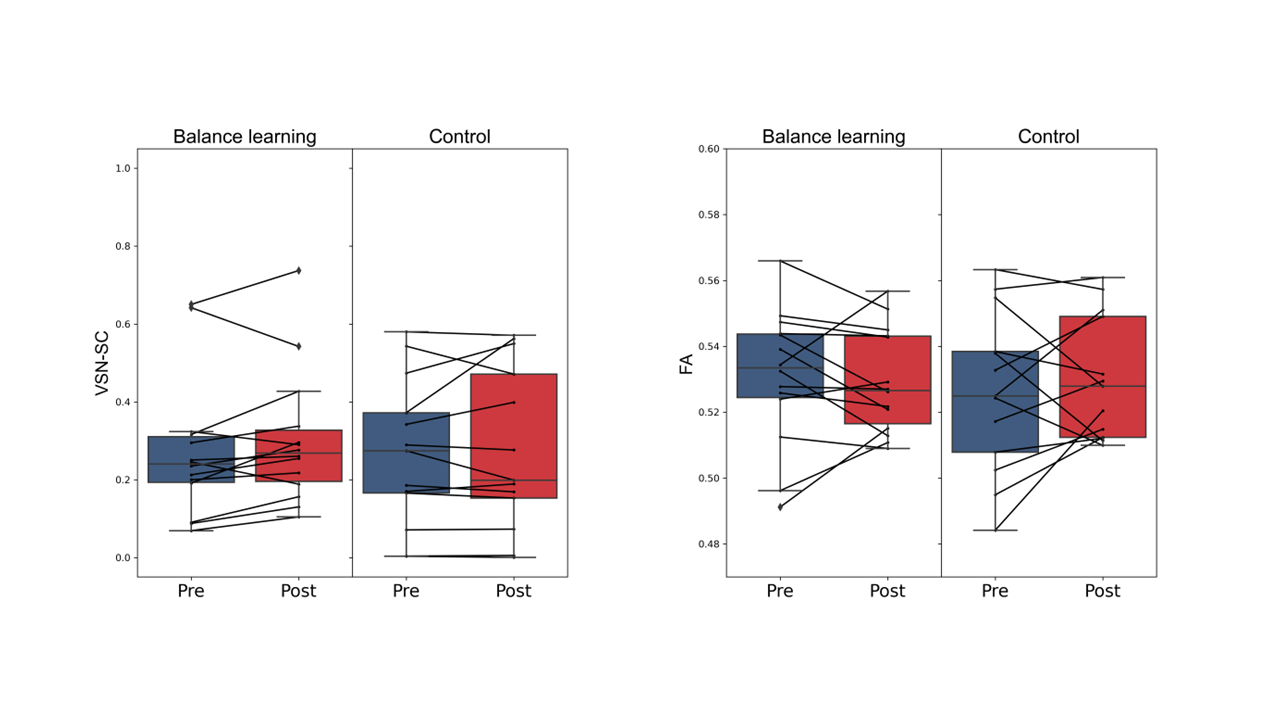

Figure 2. Comparison of ventral sensorimotor network structural connectivity (VSN-SC) and fractional anisotropy (FA) between groups. No significant interaction effect was found for the two parameters.

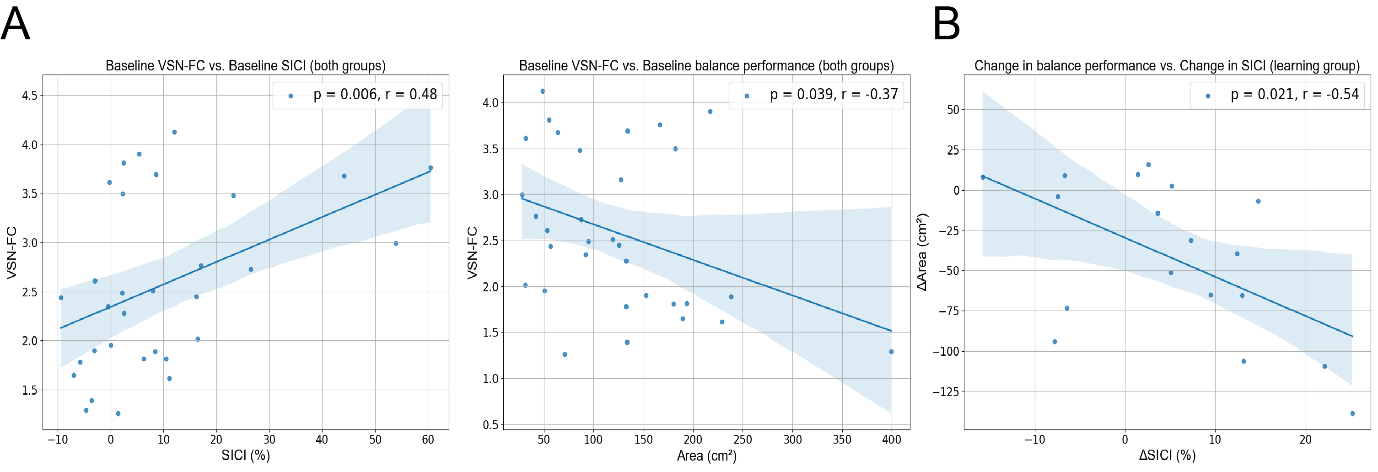

Figure 3. Replicate the correlation analysis of Figure 5 of main text but with age and sex as covariates. The results are generally consistent with what was reported in the main text (p-values are uncorrected).

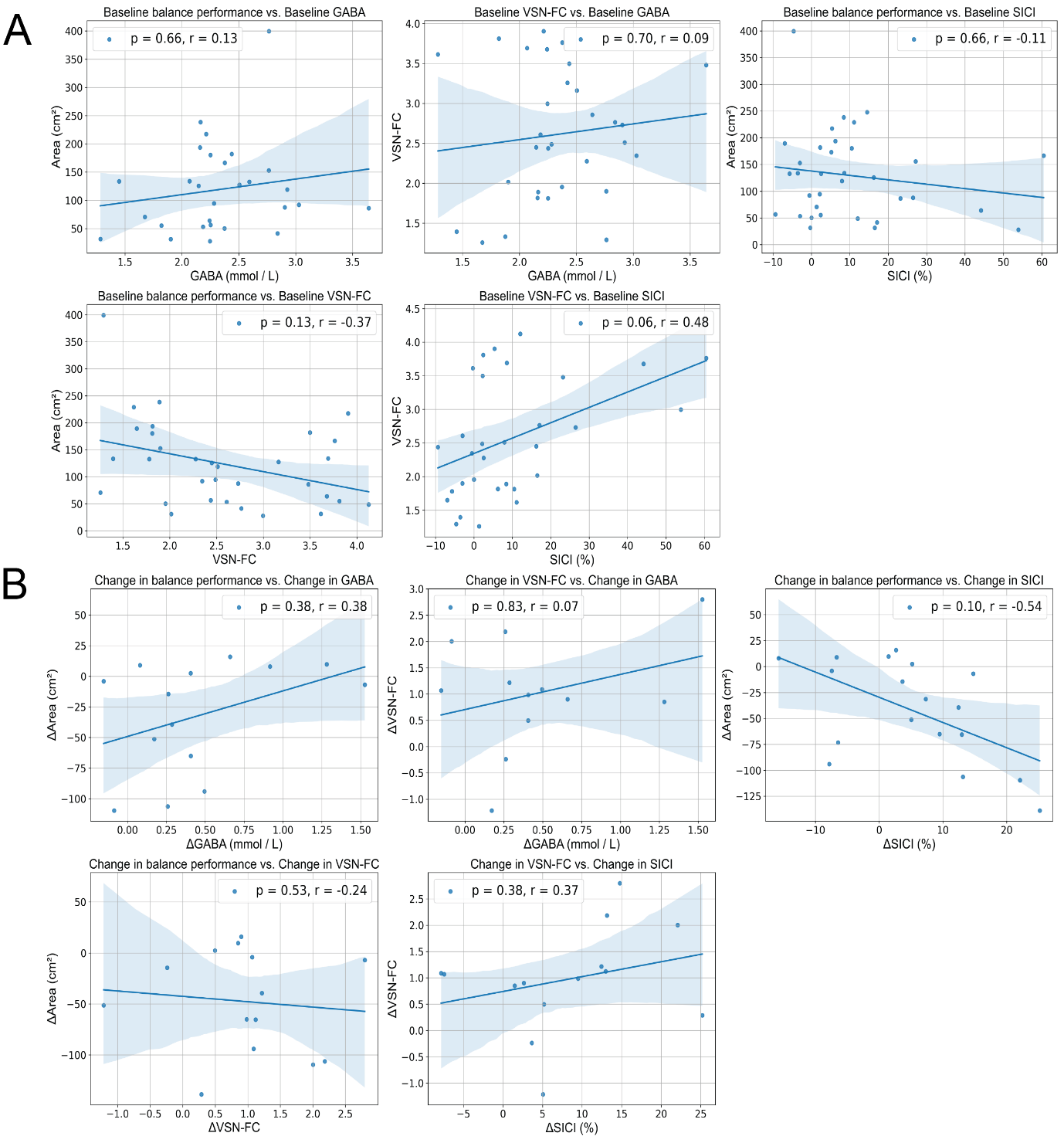

Figure 4. Correlations plots for GABA/SICI and SNFC/balance performance at baseline (A) and for delta values in learning group (B). P-values are corrected.

Table 1. Number of participants in each group for each analysis.

|  | Balance learning | | Control | |
| --- | --- | --- | --- | --- |
|  | Pre | Post | Pre | Post |
| MRS | 15 | 14 | 16 | 13 |
| fMRI | 17 | 14 | 18 | 16 |
| Balance Performance | 18 | 18 | 18 | 18 |
| SICI | 16 | 16 | 16 | 16 |

Table 2. Number of participants entered into correlation analysis. Correlation analysis was only performed when data from both time points and both measurements were available.

| Analysis | Group | Number of participants |
| --- | --- | --- |
| Baseline Performance - GABA | Both | 28 |
| Baseline VSN-FC - GABA | Both | 31 |
| Baseline Performance - SICI | Both | 34 |
| Baseline Performance – VSN-FC | Both | 32 |
| Baseline VSN-FC - SICI | Both | 31 |
| Change Performance - GABA | Learning | 14 |
| Change VSN-FC - GABA | Learning | 12 |
| Change Performance – VSN-FC | Learning | 18 |
| Change Performance - SICI | Learning | 14 |
| Change VSN-FC - SICI | Learning | 14 |

Table 3. Metabolite profiling measured using sSPECIAL.

|  |  |  | Balance learning | | | | | | Control | | | | | |
| --- | --- | --- | --- | --- | --- | --- | --- | --- | --- | --- | --- | --- | --- | --- |
|  |  |  | Pre | | | Post | | | Pre | | | Post | | |
| Metabolite | T2 relaxation time | reference | Mean Concentration (I. U.) | SD | Mean CRLB (%) | Mean Concentration (I. U.) | SD | Mean CRLB (%) | Mean Concentration (I. U.) | SD | Mean CRLB (%) | Mean Concentration (I. U.) | SD | Mean CRLB (%) |
| Asp | 97 | Marjańska M, et al. | 2.17 | 0.74 | 14.44 | 2.40 | 0.69 | 10.47 | 2.06 | 0.50 | 10.44 | 2.08 | 0.36 | 9.20 |
| Gln | 98 | Marjańska M, et al. | 0.89 | 0.35 | 13.33 | 1.06 | 0.33 | 10.27 | 0.94 | 0.22 | 9.88 | 1.01 | 0.19 | 8.93 |
| Glu | 98 | Marjańska M, et al. | 5.96 | 0.54 | 2.11 | 6.25 | 0.61 | 1.93 | 5.57 | 0.46 | 1.94 | 5.77 | 0.55 | 1.93 |
| GSH | 97 | Marjańska M, et al. | 1.10 | 0.23 | 5.39 | 1.09 | 0.26 | 5.53 | 0.97 | 0.18 | 5.38 | 1.02 | 0.12 | 4.73 |
| Ins | 100 | Marjańska M, et al. | 5.04 | 0.63 | 2.00 | 5.06 | 0.85 | 2.07 | 4.86 | 0.64 | 2.00 | 4.99 | 0.46 | 1.80 |
| Lac | 94 | Dehghani M, et al. | 0.87 | 0.52 | 8.58 | 0.72 | 0.42 | 8.93 | 0.69 | 0.35 | 9.81 | 0.74 | 0.30 | 6.21 |
| NAA | 110 | Marjańska M, et al. | 8.22 | 0.71 | 1.17 | 8.10 | 0.86 | 1.27 | 7.79 | 0.60 | 1.06 | 7.69 | 0.66 | 1.13 |
| Tau | 90 | Marjańska M, et al. | 1.09 | 0.30 | 9.56 | 1.07 | 0.22 | 8.87 | 1.13 | 0.26 | 7.00 | 1.00 | 0.27 | 8.27 |
| NAAG | 110 | Marjańska M, et al. | 1.83 | 0.46 | 4.83 | 1.90 | 0.43 | 4.47 | 1.94 | 0.44 | 3.63 | 2.07 | 0.37 | 3.47 |
| tCho | 139 | Marjańska M, et al. | 1.18 | 0.18 | 2.94 | 1.20 | 0.13 | 2.73 | 1.12 | 0.20 | 2.50 | 1.19 | 0.17 | 2.20 |
| tCr | 108 | Marjańska M, et al. | 6.26 | 0.41 | 1.39 | 6.46 | 0.59 | 1.33 | 5.77 | 0.42 | 1.25 | 5.94 | 0.45 | 1.20 |

**References**: Marjańska M, Auerbach EJ, Valabrègue R, Van de Moortele PF, Adriany G, Garwood M. Localized 1H NMR spectroscopy in different regions of human brain in vivo at 7T: T 2 relaxation times and concentrations of cerebral metabolites. NMR Biomed. 2012;25:332–339 doi: 10.1002/nbm.1754.

Dehghani M, Do KQ, Magistretti P, Xin L. Lactate measurement by neurochemical profiling in the dorsolateral prefrontal cortex at 7T: accuracy, precision, and relaxation times. Magn. Reson. Med. 2020;83:1895–1908 doi: 10.1002/mrm.28066.
